# Chemical proteomics reveals regulation of bile salt hydrolases via oxidative post-translational modifications

**DOI:** 10.64898/2025.12.11.693737

**Authors:** Amy K. Bracken, Kien P. Malarney, Pamela V. Chang

## Abstract

The gut microbiome is the vast, diverse ecosystem of microorganisms that inhabits the human intestines and provides numerous essential functions for the host. One such key role is the metabolism of primary bile acids that are biosynthesized in the host liver into a plethora of secondary bile acids produced by gut bacteria. These metabolites serve as both antimicrobial and chemical signaling agents within the host. The critical microbial enzyme that plays a gatekeeping role in secondary bile acid metabolism is bile salt hydrolase (BSH), a cysteine hydrolase that is primarily known for its deconjugating and reconjugating activities on bile acid substrates. Despite the crucial nature of these biotransformations, regulation of BSH activity is not well understood. Here, we found that the catalytic cysteine 2 (Cys2) within the BSH active site exists in multiple sulfur oxidation states including sulfenic acid (Cys-SOH). Importantly, we show this reversible oxidative post-translational modification (oxPTM) ablates BSH catalytic activity. We have leveraged this discovery to develop a chemoproteomic platform featuring a sulfenic acid-reactive bile acid probe to profile BSH Cys2 oxPTMs throughout the gut microbiome. Our results reveal that though most gut microbiota-associated BSHs exist in the active Cys2-SH state, some are preferentially and reversibly inactivated in the Cys2-SOH state. This reversible oxidation of Cys2 may serve as a general mechanism to regulate BSH activity *in vivo* in response to a changing physiological environment.

## Main

The gut microbiome is the ecosystem of trillions of microbes, including bacteria, viruses, fungi, parasites and archaea, that inhabit the intestines^1,2^. These microorganisms rival the number of cells in the human body and perform many essential functions, including digestion of dietary fiber, production of essential nutrients, and regulation of the host immune system^3^. The majority of these microbes include bacteria that are capable of producing a multitude of small-molecule metabolites that are biosynthesized by enzymes encoded by their metagenome, which contains 150-fold more genes than the human genome^4^. The resultant microbial metabolome comprises bacterial natural products such as lipids, carbohydrates, peptides, polyketides, terpenoids, and amino acid metabolites, some of which are bioactive and have important effects on myriad host physiological processes.

Bile acids are cholesterol-derived metabolites produced in the liver that are then extensively modified by numerous enzymes expressed by the gut microbiota when they are postprandially secreted into the intestines^5–7^. These molecules have multiple purposes in the body, serving as emulsifying agents to help absorb lipophilic dietary nutrients, antimicrobial agents, and signaling molecules that activate numerous host receptors. The biological impact of these metabolites is extensive as they regulate many physiological processes such as metabolism, immunity, and tumorigenesis^6,8^.

Despite the importance of these molecules, understanding the biosynthesis and biological functions of this disparate pool of bile acids is an ongoing area of research due to the diversity of their chemical structures^9,10^. Recent discovery of novel bile acids produced by the gut microbiota, termed microbial conjugated bile acids (MCBAs), has multiplied the estimated number of these metabolites from hundreds to thousands, thus considerably expanding the potential biological roles of these distinct molecules. To date, these MCBAs include bile acid amidates that contain proteinogenic amino acids, polyamines, and cysteamine conjugates^10–14^.

Central to the metabolism of bile acids is a critical gate-keeping microbial enzyme called bile salt hydrolase (BSH, EC 3.5.1.24) that is involved in many physiological processes^15,16^. This cysteine hydrolase is from the N-terminal nucleophile (Ntn) hydrolase superfamily of enzymes that canonically cleaves the C24 amide bond within bile acid conjugates produced by the liver such as glyco- and taurocholic acid (**Figure 1A**)^17^. Critically, this step is thought to precede all subsequent biotransformations carried out by the gut microbiota, including oxidation, epimerization, and 7*α*-dehydroxylation that ultimately convert primary bile acids, cholic and chenodeoxycholic acid, into secondary bile acids, deoxycholic and lithocholic acid^18^. Recent work has uncovered that BSH also possesses catalytic function as an amine *N*-acyl transferase that is responsible for the production of the newly discovered MCBAs (**Figure 1A**)^12,14^. Additionally, we and others have found that BSH can deconjugate these MCBAs, which significantly increases the complexity of bile acid metabolism carried out by the gut microbiota^19,20^.

**Figure 1.**
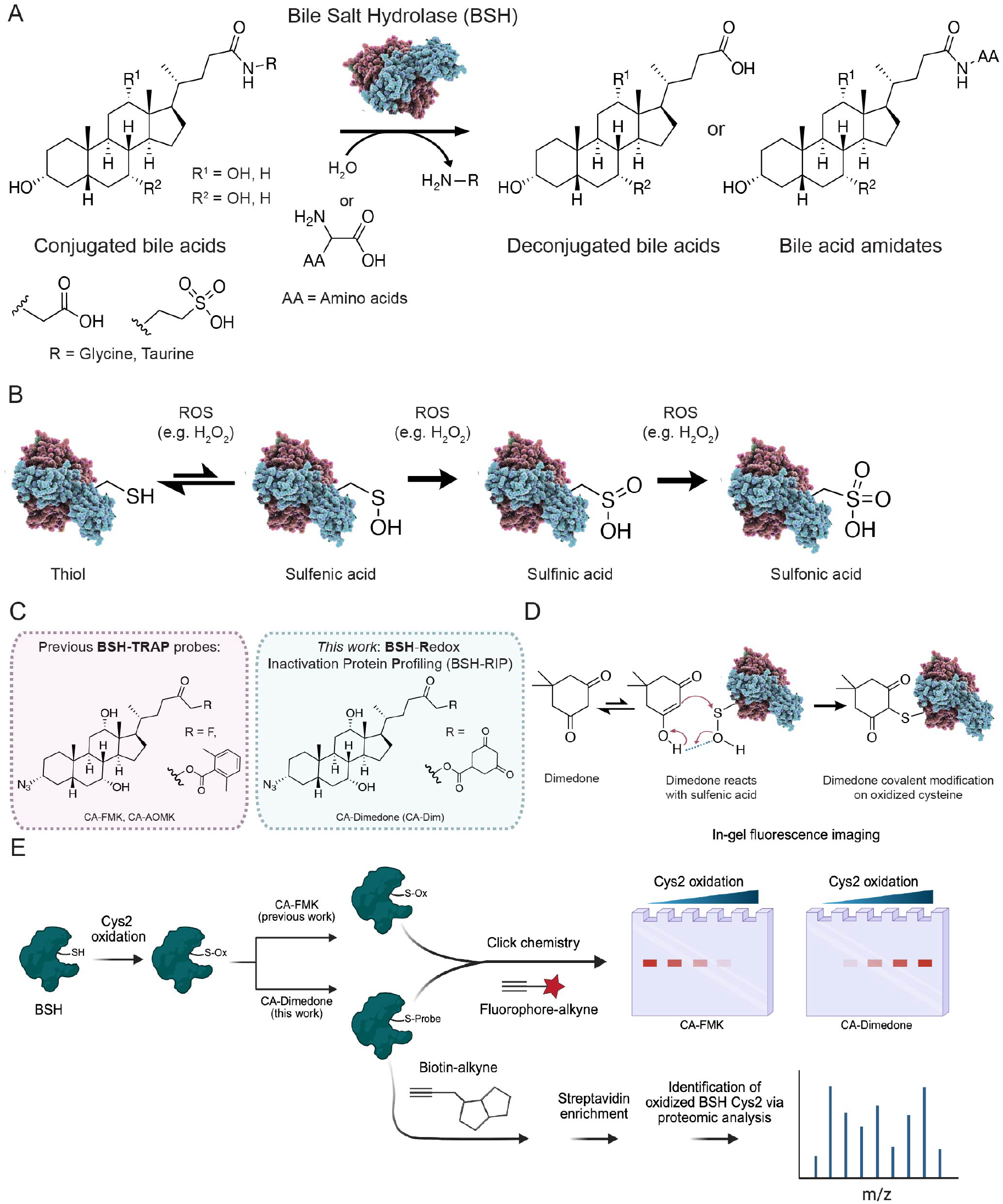
Chemoproteomic approach for profiling redox regulation of bile salt hydrolase (BSH) activity in the gut microbiome. (A) BSH controls the bile acid metabolite pool by deconjugation of conjugated bile acids (e.g., glyco- and tauro-) and N-acyl transfer of alternative amino acids onto these substrates. (B) Cysteine oxidation to higher sulfur oxidation states is a post-translational modification (PTM) that is found in proteins. (C) Previous BSH activity-based probes (ABPs) have been reported in our strategy **BSH**-**T**agging and **R**etrieval with **A**ctivity-based **P**robes (BSH-TRAP). In this work, we report the chemoproteomic platform **BSH**-**R**edox **I**nactivation protein **P**rofiling (BSH-RIP) using cholic acid-containing dimedone (CA-Dimedone), that targets inactive BSHs. (D) Dimedone acts as a nucleophile to react selectively with cysteine-sulfenic acids. (E) Overall chemical strategy, termed BSH-RIP, to probe oxidation states of BSH catalytic Cys2 residues using CA-Dimedone labeling, followed by Cu-catalyzed azide-alkyne cycloaddition (CuAAC) to tag labeled enzymes with a fluorophore or affinity handle for detection of BSH activity via in-gel fluorescence or mass spectrometry-based proteomics, respectively.

Post-translational modifications (PTMs), including phosphorylation, acetylation, and methylation, are dynamic modifications to proteins that regulate their activities in response to changes in environmental stimuli^21^. These chemical modifications are typically installed or removed by specific enzymes, called writers and erasers, and often reflect the cellular metabolic state due to the production of small-molecule metabolites in response to these environmental changes^22^.

Oxidative molecules such as reactive oxygen, nitrogen, and sulfur species produced during non-pathological conditions act as targeted second messengers to regulate biological activity via site-specific PTM of proteins^23^. Sulfur is unique in its ability to exist in multiple oxidation states (-2 to +6), so redox reactivity of cysteine residues can lead to an array of oxidative PTMs (oxPTMs), including reversible sulfenic and irreversible sulfinic and sulfonic acids (**Figure 1B**)^24,25^. An emerging paradigm suggests that reversible oxPTMs of cysteine thiols such as sulfenic acid function as binary switches to regulate protein function, activity, localization, and protein-protein interactions^26^. These Cys oxPTMs have been profiled in bacterial and mammalian cells using various chemoproteomic platforms^27–31^.

Despite the pivotal role of BSH in bile acid metabolism and important physiological processes, essentially little is known regarding the molecular mechanisms that regulate its activation^15,16^. The CGH family is categorized into two clusters based on the presence of a pre-peptide that is autocatalytically cleaved to produce a mature and active form of the enzyme^32^. However, given the sheer number of BSHs expressed by approximately 25% of human gut bacteria, and our increased understanding of BSH activity as a master regulator of bile acid metabolism, additional mechanisms beyond post-translational processing likely exist to control its activity^33^. Here, we find that BSH activity is regulated by oxPTMs of its catalytic Cys2 residue that is essential for its activity.

To characterize these Cys2 oxPTMs that regulate BSH activity within the gut microbiota on the systems biochemistry level, we have developed a chemoproteomic approach that we term **BSH**-**R**edox **I**nactivation Protein **P**rofiling (BSH-RIP). Here, we employ a chemical probe azido**C**holic **A**cid-containing **Dim**edone (CA-Dim, **Figure 1C-D**, Supporting Information, SI Scheme 1) that selectively labels inactive BSHs that harbor a Cys2 sulfenic acid (Cys-SOH) because it is based on the sulfenic acid-selective 5,5-dimethyl-1,3-cyclohexanedione (dimedone) warhead^34–36^. The Cys2 oxPTM in BSH can then be profiled using copper(I)-catalyzed azide-alkyne cycloaddition (CuAAC) tagging with either a fluorophore-alkyne or biotin-alkyne for visualization or identification, respectively (**Figure 1E**). CA-Dim was synthesized analogously to our previously developed BSH activity-based probes (ABPs)^37–39^.

To demonstrate that BSH activity is regulated by oxPTMs of the active site Cys2 residue, we first cloned and expressed BSH from *Lactiplantibacillus plantarum* and CGH from *Clostridium perfringens* as previously described^19,39^. We found that both purified BSHs have decreased activity with increasing concentrations of H_2_O_2_ using an *in vitro* biochemical assay that measures release of glycine or taurine from the BSH substrate glycocholic and taurocholic acid, respectively (**Figure 2A, Figure S1**)^40^. These results suggest that oxidation of the Cys2 leads to decreased catalytic activity. To demonstrate that the decreased BSH activity is due to sulfenic acid formation, we employed our previously reported BSH ABP, azidocholic acid containing fluoromethyl ketone, CA-FMK, whose labeling of BSH Cys2 can be competed away with dimedone under oxidizing conditions (**Figure 2B)**. After pre-treatment of samples with dimedone, followed by CA-FMK labeling, in the presence or absence of H_2_O_2_, CuAAC tagging with AZDye 647-alkyne, and SDS-PAGE analysis, we found that BSH labeling by CA-FMK was decreased in the presence of H_2_O_2_ using in-gel fluorescence imaging (**Figure 2C, Figure S2**)^37^. These data suggest that the addition of H_2_O_2_ leads to oxidation of Cys2 to higher sulfur oxidation states, including sulfenic acid, which cannot be labeled with CA-FMK. Together, these results suggest that BSH activity is controlled by oxPTMs of the active site Cys2 residue.

**Figure 2.**
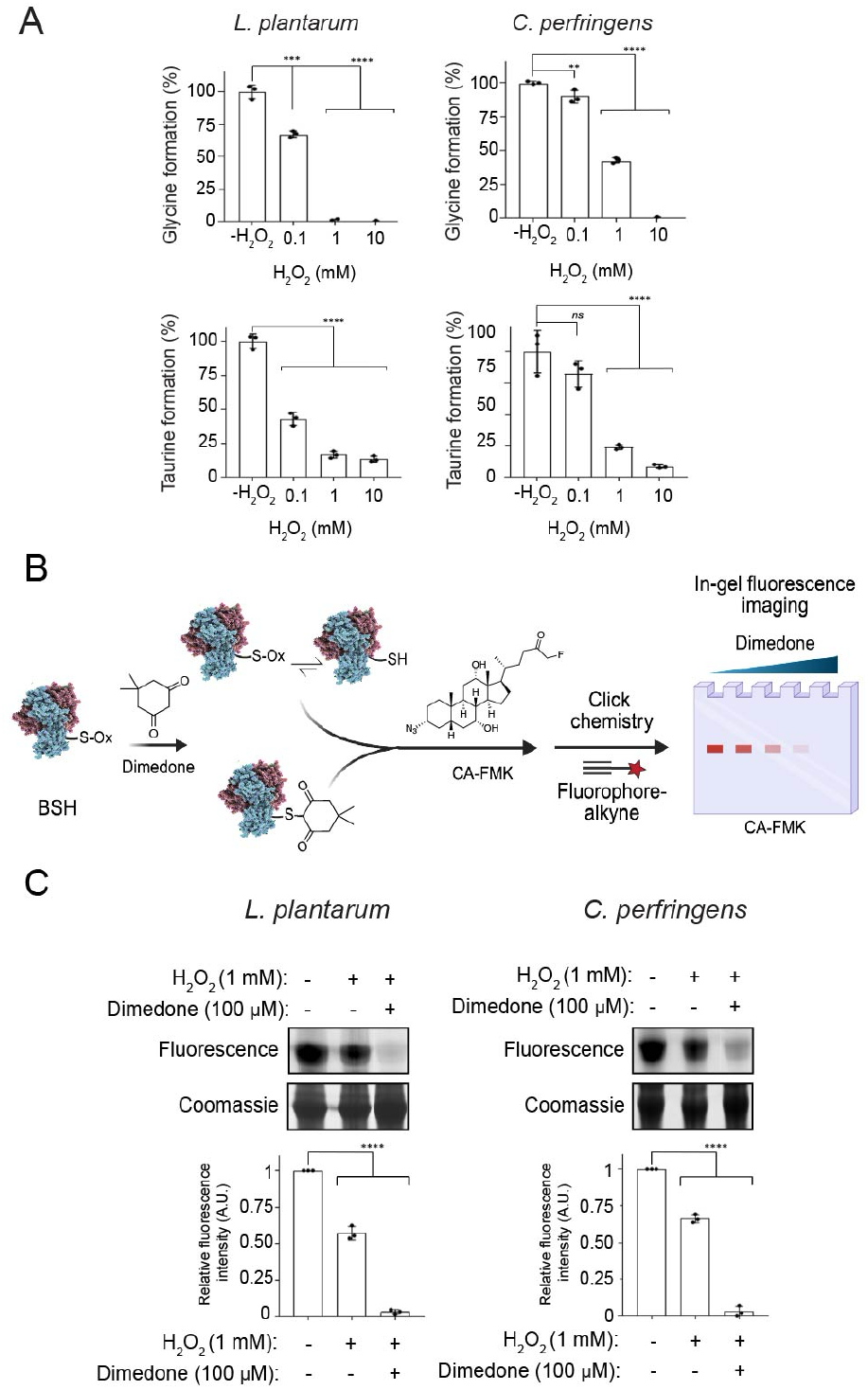
Oxidation decreases BSH activity through formation of sulfenic acid intermediates. (A) Purified *Clostridium perfringens* choloylglycine hydrolase (CGH) and *Lactiplantibacillus plantarum* BSH were treated with and without H_2_O_2_ for 1 min at the indicated concentrations prior to addition of the conjugated bile acid and incubation for 20 min at 37 °C. Percent enzyme activity was normalized relative to control. (B) Schematic for the dimedone competition experiment. (C) Purified CGH and BSH were oxidized via addition of H_2_O_2_ (1 mM) and incubated with dimedone (100 μM) for 30 min at 37 °C, followed by treatment with CA-FMK (10 μM) for 30 min at 37 °C. The samples were tagged with AZDye 647-alkyne using CuAAC and analyzed by SDS-PAGE, followed by visualization by in-gel fluorescence and Coomassie staining. The protein bands were quantified by densitometry using ImageJ (bottom panels). A.U. = arbitrary units. Error bars represent standard deviation from the mean. One-way ANOVA was performed, followed by post hoc Tukey’s test: * p<0.05, ** p< 0.01, *** p<0.001, **** p<0.0001, n.s. = not significant, n = 3.

Based on these findings, we next sought to demonstrate that *C. perfringens* BSH Cys2 can be oxidized to sulfenic acid and additional sulfur oxidation states. In these studies, we treated purified *C. perfringens* BSH with or without H_2_O_2_, followed by dimedone or vehicle, and performed peptide mapping using liquid chromatography-tandem MS (LC-MS/MS). We found that BSH-Cys2 can exist in its reduced (Cys2-SH) and oxidized states (Cys2-SOH and Cys2-SO_2_H) as determined by the expected masses of the sulfenic acid-dimedone adduct, sulfenic, and sulfinic acid modifications (**Figure 3, Figure S3, Table S1**). These data are consistent with existing reports in which *C. perfringens* BSH can be crystallized in multiple Cys2 oxidation states^41–43^. These results support our findings that the catalytic Cys2 within BSH can exist in higher sulfur oxidation states, rendering it inactive to catalysis.

**Figure 3.**
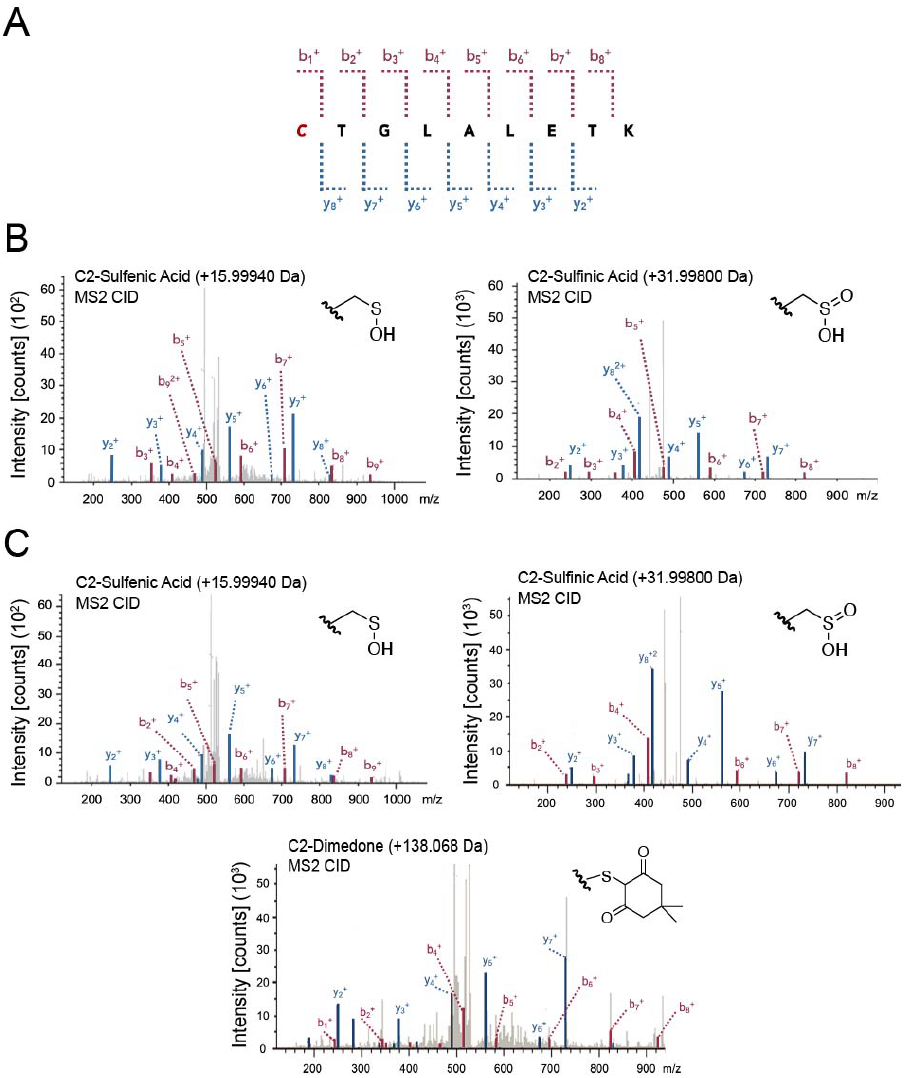
LC-MS/MS analysis of *C. perfringens* CGH reveals oxidative PTMs (oxPTMs) that occur on Cys2. *C. perfringens* CGH was incubated with dimedone (100 μM) or vehicle for 30 min prior to LC-MS/MS analysis. (A) Peptide containing *C. perfringens* Cys2 (red, italics) that was detected by LC-MS/MS analysis. Anticipated b and y ions are indicated via the dotted red and blue lines, respectively. (B) Representative MS/MS (MS^2^) spectra showing detected oxPTMs on the Cys2 peptide from CGH treated with vehicle. Both the sulfenic (Cys-SOH, *left*) and sulfinic (Cys-SO_2_H, *right*) acid states of Cys2 were present. (C) Representative MS^2^ spectra showing detected oxPTMs and covalent dimedone labeling of sulfenic acid on the Cys2 peptide from CGH treated with dimedone. Both the sulfenic (Cys-SOH, *left*) and sulfinic (Cys-SO_2_H, *right*) acid states of Cys2 were present as well as the dimedone-modified peptide containing sulfenic acid (*bottom*).

Given that BSH Cys2 can exist in its oxidized form as a sulfenic acid, we next determined that CA-Dim can label Cys2-SOH in BSHs from *L. plantarum* and *C. perfringens*. In these studies, treatment of purified *L. plantarum* BSH with dithiothreitol (DTT), a reducing agent that reduces sulfenic acid to the free thiol, decreased labeling with CA-Dim as detected by CuAAC tagging with AZDye 647-alkyne and in-gel fluorescence imaging (**Figure 4A, Figure S4A**). In contrast, DTT treatment of *L. plantarum* and *C. perfringens* BSHs did not alter CA-FMK labeling. We also found that mutation of the catalytic Cys2 to serine (Ser2) in *C. perfringens* BSH (catalytically inactive C2S mutant) led to decreased labeling with CA-Dim, as expected^19,39^. Conversely, addition of H_2_O_2_ to purified *L. plantarum* BSH led to increased labeling with CA-Dim at lower H_2_O_2_ concentrations, whereas higher concentrations of H_2_O_2_ exhibited decreased CA-Dim labeling, likely due to irreversible oxidation of Cys2 to the sulfinic and sulfonic acids (**Figure 4B, Figure S4B**). Interestingly, addition of H_2_O_2_ to purified *C. perfringens* BSH led to no change in labeling with CA-Dim. These results suggest that the *C. perfringens* Cys2 may be less solvent exposed and more difficult to oxidize than *L. plantarum* BSH Cys2. As expected, CA-Dim also did not label the *C. perfringens* C2S mutant. Collectively, these results demonstrate that CA-Dim is selective for BSH Cys2 sulfenic acid.

**Figure 4.**
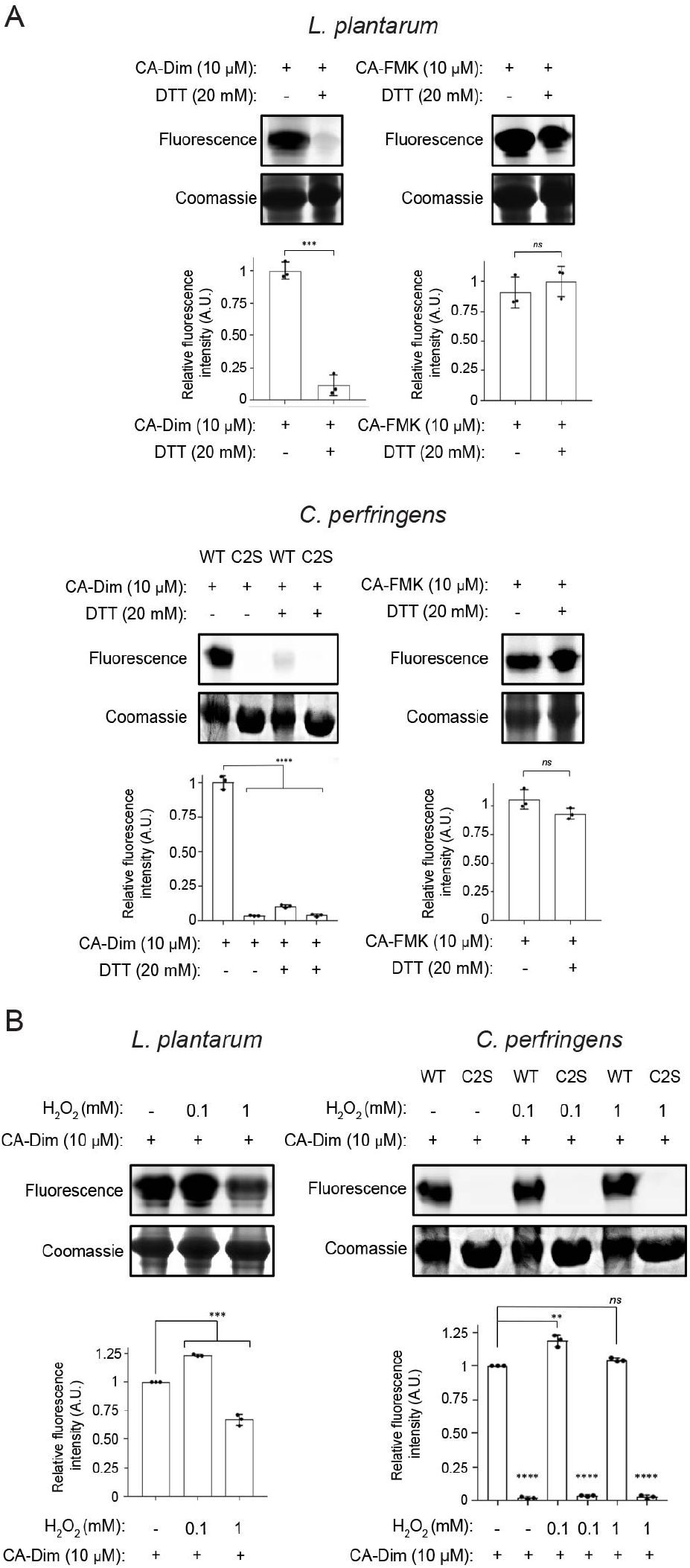
Redox state of BSH Cys2 affects labeling with CA-Dim and CA-FMK probes. (A) Purified *L. plantarum* BSH and *C. perfringens* wildtype (WT) or Cys2Ser (C2S) mutant CGH were treated with DTT (20 mM) for 15 min at 55 °C, followed by CA-Dim or CA-FMK (10 μM) for 30 min at 37 °C. (B) Purified *L. plantarum* BSH and *C. perfringens* wildtype (WT) or Cys2Ser (C2S) mutant CGH were treated with indicated concentrations of H_2_O_2_ for 1 min, followed by CA-Dim or CA-FMK (10 μM) for 30 min at 37 °C. (A-B) The samples were tagged with AZDye 647-alkyne using CuAAC and analyzed by SDS-PAGE, followed by visualization by in-gel fluorescence and Coomassie staining. The protein bands were quantified by densitometry using ImageJ (bottom panels). A.U. = arbitrary units. Error bars represent standard deviation from the mean. For comparisons between two groups, unpaired t-tests with two-tailed P values were performed: *** p<0.001, n.s. = not significant, n = 3. For comparisons between three or more groups, one-way ANOVA was performed, followed by post hoc Tukey’s test: * p<0.05, ** p< 0.01, *** p<0.001, **** p<0.0001, n.s. = not significant, n = 3.

To demonstrate the physiological relevance of these findings, we next cultured human gut bacteria *Bifidobacterium longum* subspecies *infantis* and *Bacteroides fragilis* in the presence of CA-Dim and CA-FMK. Labeling of BSHs with both CA-Dim and CA-FMK from *B. longum* and *B. fragilis* increased over time, as determined by CuAAC tagging with AZDye 647-alkyne, SDS-PAGE analysis, and in-gel fluorescence imaging (**Figure 5A-B, Figure S5A-B**). These results suggest that BSH Cys2-SH is oxidized to Cys2-SOH during culture, likely due to trace amounts of O_2_ in the anaerobic chamber, and CA-Dim is able to shift the equilibrium of this reversible reaction by covalently trapping the sulfenic acid. We also determined the specificity of sulfenic acid labeling within BSH by pre-treating live *B. longum* and *B. fragilis* with dimedone, followed by CA-Dim or CA-FMK (**Figure 5C-D, Figure S5C-D**). Dimedone was able to compete away BSH labeling by both probes, suggesting that the Cys2-SOH sulfenic acid can be covalently trapped over time in live gut anaerobes. We next determined that CA-Dim can enrich for BSH Cys2 oxPTMs in live *B. longum* and *B. fragilis* BSHs as determined by mass spectrometry (MS)-based proteomics (**Figure 5E-F, Figure S5E-F**). In these studies, we cultured live bacteria with CA-Dim, followed by CuAAC tagging with biotin-alkyne, streptavidin enrichment, and shotgun proteomics (**Figure S6, Table S2**). Collectively, these results indicate that oxPTMs of BSH Cys2 exist in human gut bacteria and that these redox states can be profiled in live gut anaerobes using our chemoproteomic platform.

**Figure 5.**
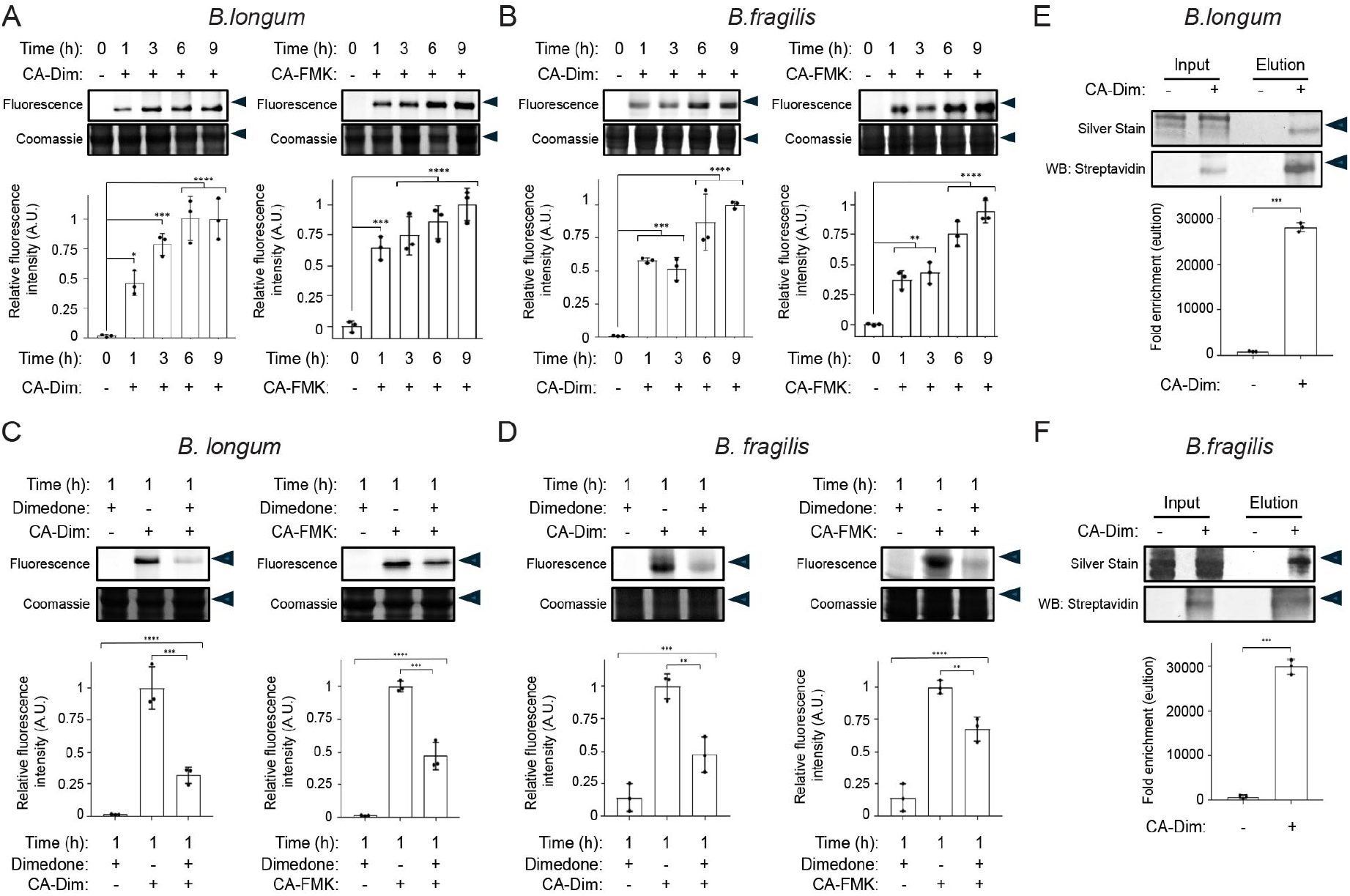
CA-Dim labels sulfenic acid in live human gut anaerobes. (A) *Bifidobacterium longum* subsp. *infantis* and (B) *Bacteroides fragilis* were treated with CA-Dim or CA-FMK (100 μM) for the indicated amount of time. (C) *B. longum* subsp. *infantis* and (D) *B. fragilis* were treated with dimedone (1 mM) for 1 h, followed by CA-Dim or CA-FMK at 100 μM for 1 h. (A-D) Bacterial lysates were tagged using CuAAC with AZDye 647-alkyne and analyzed by SDS-PAGE, followed by visualization by in-gel fluorescence and Coomassie staining. (E-F) *B. longum* and *B. fragilis* was treated with CA-Dim (100 μM) for 12 h. Bacterial lysates were tagged using CuAAC with biotin-alkyne, followed by enrichment of labeled protein via streptavidin-agarose pulldown. Samples were analyzed by SDS-PAGE, followed by silver stain or Western blot using streptavidin-horse radish peroxidase (HRP). Arrow indicates 37 kDa. The protein bands were quantified by densitometry using ImageJ (bottom panels). A.U. = arbitrary units. Error bars represent standard deviation from the mean. One-way ANOVA, followed by post hoc Tukey’s test: * p<0.05, ** p< 0.01, *** p<0.001, **** p<0.0001, n.s. = not significant, n = 3.

Next, we applied BSH-RIP to profile BSH Cys2 oxPTMs in the mouse gut microbiome. Fecal bacteria were isolated, lysed, and anaerobically labeled with CA-Dim and CA-FMK to identify changes in BSH Cys2 redox chemistry (**Figure 6A, Figure S7A**). Both probes showed increasing labeling over time, with CA-Dim labeling peaking at 1 h, likely due to reversible oxidation to the sulfenic acid. We also determined that probe labeling is concentration-dependent, consistent with covalent adduction of CA-Dim and CA-FMK to gut microbiota-associated BSHs (**Figure 6B, Figure S7B**).

**Figure 6.**
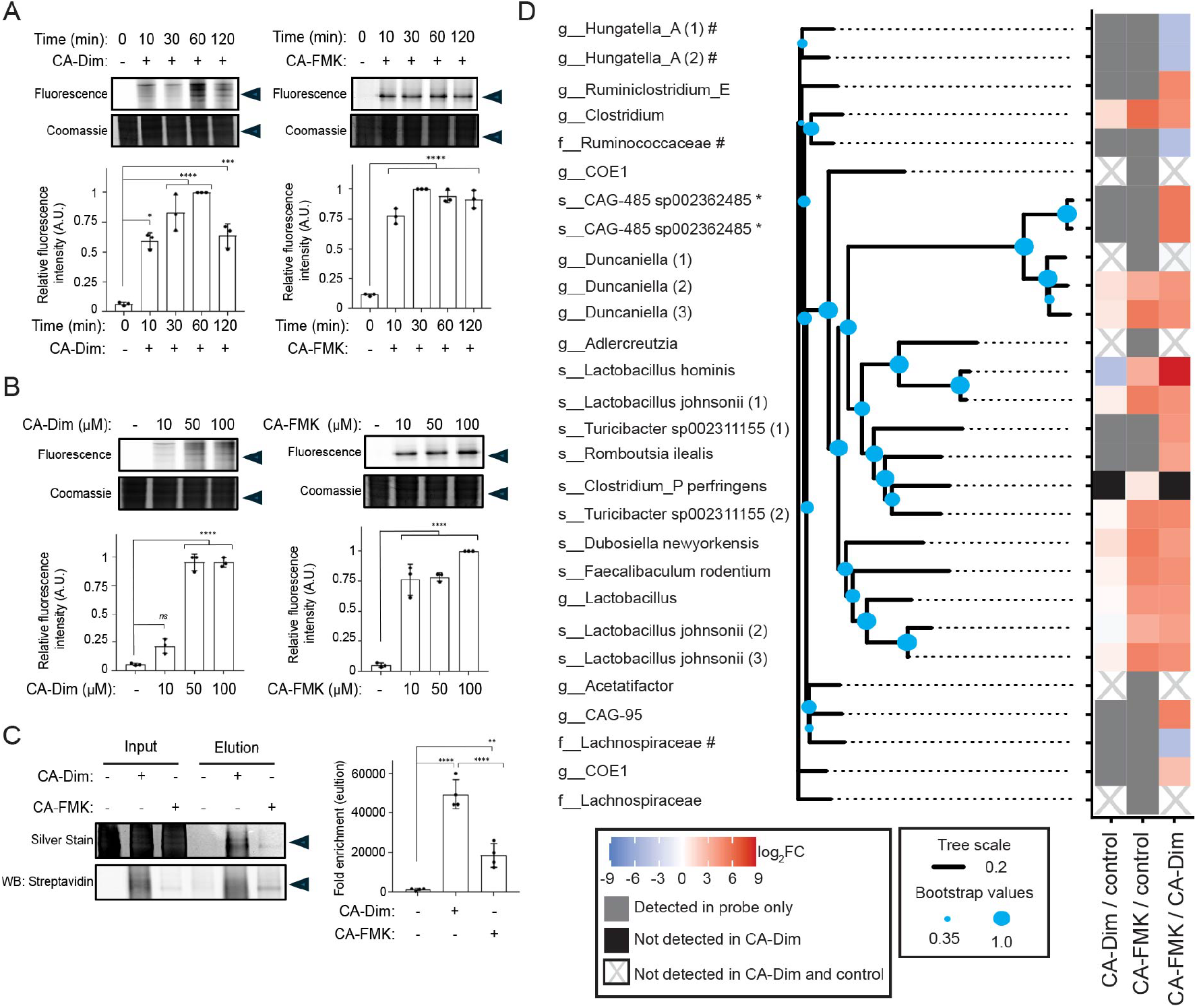
Chemoproteomic profiling of oxPTMs of BSH Cys2 in the gut microbiome. (A) Bacteria isolated from fecal samples of wildtype mice were lysed and treated with CA-Dim or CA-FMK (100 μM) for the indicated amount of time. (B) Alternatively, bacteria isolated from fecal samples of wildtype mice were lysed and treated with CA-Dim or CA-FMK at the indicated concentrations for 1 h. (A-B) The samples were tagged using CuAAC with AZDye 647-alkyne, analyzed by SDS-PAGE, and visualized by in-gel fluorescence and Coomassie Blue staining. (C) Bacteria isolated from fecal samples of wildtype mice were lysed and treated with CA-Dim or CA-FMK (100 μM) for 1 h. The samples were tagged using CuAAC with biotin-alkyne, followed by enrichment via streptavidin-agarose pulldown. Elutions were analyzed by SDS-PAGE, followed by silver stain or Western blot. (A-C) The arrow indicates the protein ladder position at 37 kDa. The gel bands were quantified by densitometry using ImageJ (bottom panels). A.U. = arbitrary units. Error bars represent standard deviation from the mean. One-way ANOVA, followed by post hoc Tukey’s test: * p<0.05, ** p< 0.01, *** p<0.001, **** p<0.0001, n.s. = not significant, n = 3 (A-B), n = 4 (C). (D) Enriched samples were analyzed by MS-based proteomics. Heatmap represents bacterial BSHs that were identified. Red indicates increased BSH labeling comparing CA-FMK to CA-Dim, shown in log2 fold change, and blue indicates decreased BSH labeling comparing CA-FMK to CA-Dim. Gray indicates labeling detected in probe only, and black indicates not detected in CA-Dim. White with gray X indicates not detected in CA-Dim and control. *, # indicates that these distinct bacteria could not be fully distinguished at the taxonomic level because of the incomplete nature of shotgun metagenomic assembly. Tree scale represents phylogenetic distance determined by BLOSUM6232 score^49^. Bootstrap confidence levels are indicated^50,51^.

We then treated fecal bacterial lysates with either CA-Dim or CA-FMK to profile BSH Cys2 oxPTMs throughout the mouse gut microbiota, followed by labeling with biotin-alkyne, streptavidin enrichment, and MS-based proteomics (**Figure 6C-D, Figure S7C-E, Tables S3-S4**). CA-FMK generally exhibited stronger enrichment of gut microbiota-associated BSHs compared to CA-Dim, suggesting that the majority of BSHs within the murine gut microbiome exist in their reduced, active Cys2-SH state (**Table S4**). However, several BSHs that were enriched did not show a preferred redox state (thiol vs. sulfenic acid), which indicates that the oxidation state of these Cys2 residues could be highly dynamic or in tight equilibrium with multiple forms. When the amino acid sequences of these BSHs were clustered based on similarity, we found a strong correlation between sequence similarity and relative reactivity with CA-FMK and CA-Dim (**Figure 6D**). These results suggest that the BSH protein sequence may influence redox regulation of its Cys2, likely via differences in electrostatics and redox potential within the active site environment^46^.

In summary, we found that an important family of bacterial gatekeeping metabolic enzymes within the gut microbiota, BSHs, are regulated via oxPTMs of their catalytic Cys2 residue. The reversible nature of this oxPTM enables it to act as a redox switch that controls BSH activity. As thousands of BSHs are broadly expressed in the gut microbiome, their individual and collective activities need to be tightly regulated to control secondary bile acid metabolism. Notably, our chemoproteomic studies revealed that regulation of BSH activity via these oxPTMs is strongly related to the BSH protein sequence, which we propose dictates key features of the active site local environment including the redox potential of Cys2. In this redox-based PTM model, each BSH enables regulation of its own activity based on environmental conditions in the gut. This model would elegantly solve a challenging problem for how a diverse and genetically complex community of microorganisms is able to rapidly adapt to environmental changes and change their metabolic output, in contrast to alternative mechanisms such as transcriptional control. Moving forward, we envision that BSH-RIP, in combination with existing activity-based protein profiling tools, will represent a useful systems biochemistry strategy for understanding redox regulation of BSH activity in different disease states associated with oxidative stress in the gut^37– 39,47,48^.

## TOC graphic

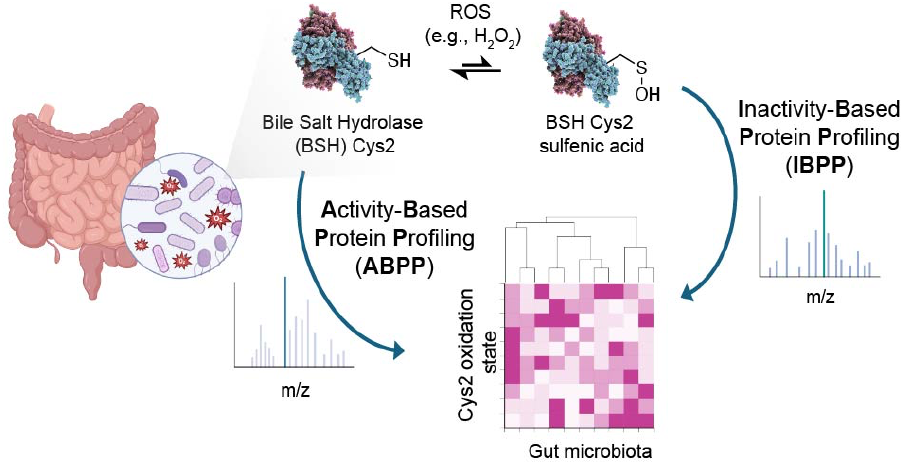

## Associated content

**Supporting Information is available free of charge at**

(PDF)

Table S1. C. *perfringens* CGH

Table S2. *B. longum* subsp. *infantis* and *B. fragilis*

Table S3. Mouse microbiome proteomics file (raw)

Table S4. Mouse microbiome proteomics file (analyzed)

## Notes

The authors declare no competing financial interests.

## Acknowledgments

This work was supported by an NIH R35 Maximizing Investigators’ Research Award for Early Stage Investigators (R35GM133501). Research in the Chang Lab is supported by a Beckman Young Investigator Award (to P.V.C.) from the Arnold and Mabel Beckman Foundation and a Sloan Research Fellowship (to P.V.C.) from the Alfred P. Sloan Foundation. A.K.B. and K.P.M. were supported by an NIH Chemistry-Biology Interface Predoctoral Training Grant (T32GM138826) and an NSF GRFP (to A.K.B. and K.P.M.). This work made use of the Cornell University NMR Facility, which is supported, in part, by the NSF through MRI award CHE-1531632. We are grateful to Samantha Scott for animal husbandry and to Sheng Zhang and Qin Fu for helpful discussions. We also thank the Weill Institute for Cell and Molecular Biology for additional resources.

## Author Information

### Author

Amy K. Bracken-Department of Chemistry and Chemical Biology, Cornell University, 930 Campus Road, Ithaca, NY 14853, United States;

Kien P. Malarney-Department of Microbiology, Cornell University, 930 Campus Road, Ithaca, NY 14853, United States;

Complete contact information is available at:

## Notes

### Competing Interest Statement

The authors have declared no competing interest.

