## Supporting Information for "Chemical proteomics reveals regulation of bile salt hydrolases via oxidative post-translational modifications"

Identification of Enriched BSHs from Mouse Fecal Bacteria using LC-MS/MS Analysis: 10

### Biological Methods

**General Materials:**

UV absorbance measurements were acquired on a Bio-Tek Synergy H1 Hybrid Multi-mode R microplate spectrophotometer. The anaerobic chamber (Coy Lab, model AC16-113) was maintained under the gas composition: 3% hydrogen, 20% carbon dioxide, and 77% nitrogen. Coy Lab (model 2000) incubator was used inside the anaerobic chamber for the growth of anaerobic bacteria.

**Bacterial Cultures:**

*Bifidobacterium longum* subsp. *infantis* ATCC 15697 and *Bacteroides fragilis* ATCC 25285 were grown anaerobically in Bifidobacterium broth and BHI broth, respectively, at 37 °C until reaching stationary phase, after which they were aliquoted into microfuge tubes, pelleted (4,500 x g, 15 min, 4°C), flash frozen, and stored at -80 °C until further use.

***C. perfringens* CGH and *L. plantarum* BSH Protein Expression and Purification:**

The previously reported[1,2] pET-21b expression plasmid containing *C. perfringens* wildtype (WT) or C2S CGH or *L. plantarum* WT BSH was introduced into *Escherichia coli* Rosetta BL21 competent cells and grown overnight on LB agar plates with ampicillin (50 µg/mL). Resulting colonies were picked and cultured in Terrific Broth (12 g/L Bacto Tryptone, 23.9 g/L yeast extract, 8 ml/L glycerol, 0.22 g/L $KH_2PO_4$, 0.94 g/L $K_2HPO_4$) with ampicillin (50 µg/ml) at 37 °C to an OD600 of 0.5-0.6. CGH expression was induced by the addition of IPTG (0.1 mM), and the bacteria were incubated with shaking (220 rpm) at 18 °C for 18 h. Bacteria were then harvested via centrifugation at 4,596 x g for 30 min at 4°C. The supernatant was discarded, then the pellet was resuspended in 1X PBS prior to lysis by probe sonication (10 x 30 sec on/30 sec off). The tube was centrifuged at 4,596 x g for 30 min at 4 °C, and the supernatant was collected and incubated with Ni-NTA resin for 1.5 h at 4 °C with gentle rocking. The resin was thoroughly washed with 1X PBS containing 0.25 mM TCEP and 20 mM imidazole. Then, proteins were eluted using 1X PBS containing 0.25 mM TCEP and 500 mM imidazole (3 x 500 µL). Elutions were concentrated, and buffer exchanged into 1X PBS using Amicon ultracentrifuge concentrators (3 kDa cutoff). The purified protein was quantified by the DC assay (Bio-Rad), aliquoted into 100 µg fractions, flash frozen, and stored at -80 °C until further use.

**In-gel Fluorescence Assays with Purified CGH and BSH:**

Purified BSH (15 µg) was added to labeling buffer (50 mM NaOAc, pH 5.5 in 1X PBS) to a final volume of 94 µL. Then, vehicle (dd$H_2O$) or 100 mM $H_2O_2$ was added to the mixture for 1 min prior to the addition of CA-FMK (1 mM, 5 µL), CA-Dimedone, or DMSO. For DTT reduction experiments, DTT (20 mM) was added to each sample, which were then incubated at 55 °C for 15 min prior to addition of vehicle or probe. After incubating the reactions for 30 min at 37 °C, 200 µg of BSA was added to each sample, which were then quenched via protein precipitation using 200 µL MeOH, 75 µL of $CHCl_3$, and 100 µL of water. Samples were centrifuged at 17,000 x g for 10 min, washed with 1 mL of MeOH, and centrifuged again at 17,000 x g for 10 min. After drying protein pellets for 10 min at 37 °C, they were resuspended in 95 µL click buffer (0.1 M sodium phosphate, 4% SDS, pH 7.4) via vortexing. Reagents for Cu(I)-catalyzed azide-alkyne cycloaddition (CuAAC) were added into a master mix in the following final concentrations: 2 mM THPTA, 1 mM $CuSO_4$, 10 µM AZDye 647-alkyne (Click Chemistry Tools), and 50 mM sodium ascorbate, in a final reaction volume of 120 µL. Click reactions were performed for 1 h at 37 °C, then quenched via protein precipitation as previously described. Pellets were resuspended in 30 µL of 1X Laemmli buffer and boiled at 95 °C for 10 min, then analyzed by SDS-PAGE. Gels were destained (40% methanol, 50% acetic acid, 10% dd$H_2O$) on a rocker in the dark for 20 min, then visualized using a Bio-Rad ChemiDoc MP Imaging System. Lastly, gels were stained with Coomassie Brilliant Blue dye for 2 min, then destained (30% methanol, 5% acetic acid, 65% dd$H_2O$) overnight with gentle rocking prior to visualization using a Bio-Rad ChemiDoc MP Imaging System.

**Dimedone Competition Fluorescence Assays with Purified CGH and BSH:**

Purified BSH (15 µg) was added to labeling buffer (50 mM NaOAc, pH 5.5 in 1X PBS) to a final volume of 94 µL. Then, vehicle (dd$H_2O$) or 100 mM $H_2O_2$ was added to the mixture for 1 min prior to the addition of 1 mM dimedone (5 µL). After incubating the reactions for 30 min at 37 °C, CA-FMK (10 µM) was added to each sample, which were left to react for an additional 30 min at 37 °C. Then, BSA (200 µg) was added to each sample, which were then quenched via protein precipitation as described above. Subsequent click chemistry and fluorescence analysis was carried out as described in the section above, "In-gel Fluorescence Assays with Purified CGH and BSH."

**Activity Assays with Purified CGH and BSH:**

Assays were performed as described previously with some modification.[3] Purified BSH (60 ng) was added to assay buffer (50 mM NaOAc, pH 5.5, 107 mM $\beta$-mercaptoethanol, 55 mM EDTA), and then $H_2O_2$ or vehicle (dd$H_2O$) were added at the indicated concentrations. After 1 min incubation, respective substrate (36 mM glycocholate or taurocholate) was added, and reactions were incubated for 20 min at 37 °C. Trichloroacetic acid (50 µL) was added to quench the reaction, samples were centrifuged at 17,000 x g for 1 min to remove precipitate, and 20 µL of supernatant was removed for subsequent steps. Color development solution (4% w/v ninhydrin in 2-methoxyethanol, 0.16% w/v $SnCl_2$ in 200 mM citrate buffer, pH 5) was added to the supernatant to a total volume of 220 µL, and samples were boiled at 95 °C for 20 min. For final absorbance measurements, 122 µL of each sample was removed and added to a 96-well plate, and OD600 values were obtained for each.

**In-gel Fluorescence Assays with Live Labeling of Anaerobes:**

*B. longum* and *B. fragilis* were anaerobically grown as described above in 1 mL broth at 37 °C until the stationary phase was reached. Then, CA-FMK (20 mM, 5 µL), CA-Dimedone probe, or DMSO was added. Bacteria were grown for the indicated times, then were harvested by pelleting (4,500 x g, 15 min, 4 °C). Samples were flash frozen and stored at -80 °C until further use.

Bacterial pellets were washed 2x with cold 1X PBS, then resuspended in lysis buffer (1X PBS, 1 mM PMSF, 100 µL final volume). The samples were then transferred to a 2 mL screwcap tube. Zirconia/silica beads (110 mg) were added to each tube, which were then subjected to agitation on a bead beater at 2,400 rpm (6 x 1 min, with 3 min rest on ice between). The samples were centrifuged at 17,000 x g for 15 min to remove the beads and cell debris. The supernatant for each sample was transferred into a 1.5 mL microfuge tube, and the protein concentrations were measured using the DC assay (Bio-Rad).

For CuAAC tagging, bacterial lysates (100 µg) were diluted to 95 µL using 1X PBS. Reagents were added into a master mix in the following final concentrations: 2 mM THPTA, 1 mM $CuSO_4$, 10 µM AZDye 647-alkyne, and 50 mM sodium ascorbate, in a final reaction volume of 120 µL. Click reactions were performed for 1 h at 37 °C then quenched via protein precipitation as previously described. Pellets were resuspended in 30 µL of 1X Laemmli buffer and boiled at 95 °C for 10 min, then analyzed by SDS-PAGE. Gels were destained (40% methanol, 50% acetic acid, 10% $ddH_2O$) on a rocker in the dark for 20 min, then visualized using a Bio-Rad ChemiDoc MP Imaging System. Lastly, gels were stained with Coomassie Brilliant Blue dye for 2 min, then destained (30% methanol, 5% acetic acid, 65% $ddH_2O$) overnight with gentle rocking prior to visualization using a Bio-Rad ChemiDoc MP Imaging System.

**Dimedone Competition Fluorescence Assays in Live Anaerobes:**

*B. longum* and *B. fragilis* were anaerobically grown as described above in 1 mL broth anaerobically at 37 °C until the stationary phase was reached. Then, dimedone (100 mM, 10 µL) or vehicle (DMSO) was added to each sample for 1 h at 37 °C. Following treatment, bacteria were quickly pelleted, washed with 1X PBS, and resuspended in 1 mL of the corresponding media. Then, CA-FMK (20 mM, 5 µL), CA-Dimedone probe, or DMSO was added. Bacteria were treated for 1 h at 37 °C, then were harvested by pelleting (4,500 x g, 15 min, 4 °C). Samples were flash frozen and stored at -80 °C until further use. Lysate preparation and subsequent click chemistry and fluorescence analysis were carried out as described in the section above, "In-gel Fluorescence Assays with Live Labeling of Anaerobes."

**Propidium Iodide Staining of Live Labeled Anaerobes:**

The protocol was performed as described by the manufacturer, with some minor modifications. After treatment in an anaerobic chamber with CA-Dim or CA-FMK (100 µM) for 9 h, or dimedone (1 mM) plus CA-Dim and CA-FMK (100 µM) for 1 h each, respectively, bacteria were spun down (10,000 x g, 5 min) and washed three times with degassed, sterile 1X PBS. Each bacterial sample was then diluted to OD600 0.1, and supernatant (100 µL) was removed for subsequent steps. Then, propidium iodide (final concentration 8 ng/µL) was added to each sample, which was then incubated in the dark for 20 min. Samples were kept on ice prior to analysis via flow cytometry using the Cy3 channel on a BD Accuri C6 Plus Flow Cytometer.

**Biotin-Streptavidin Enrichment of Labeled Protein from Live Anaerobes:**

Live bacteria were treated with probe (100 µM) or vehicle (DMSO) as described above for 12 h. Lysates (1 mg) were then prepared as described above and diluted to a final volume of 500 µL with 0.2% SDS for preclearing. Streptavidin agarose beads (10 µL per 1 mg of lysate) were washed three times with 1 mL of 1X PBS containing 0.2% SDS, then added to the samples and incubated for 1 h with rotation. After preclearing, the samples were precipitated, dried for 10 min at 37°C, and resolubilized in click buffer (0.2% SDS final concentration, 1X PBS, pH 7.4). Reagents were added into a master mix in the following final concentrations: 2 mM THPTA, 1 mM $CuSO_4$, 100 µM biotin-PEG4-alkyne (biotin-alkyne, Click Chemistry Tools), and 50 mM sodium ascorbate, in a final reaction volume of 600 µL. Click reactions were performed for 1 h at 37 °C, then quenched via protein precipitation. After drying the pellets for 10 min at 37 °C, they were resuspended in PBS with a final concentration of 0.2% SDS. Streptavidin agarose beads (10 µL per 1 mg lysate) were washed three times with 1 mL of 0.2% SDS in 1X PBS, then added to the samples and incubated for 1 h with rotation. After enrichment, samples were centrifuged at 2,500 x g for 1 min at 4 °C, the supernatant was discarded, and the beads were then washed sequentially (3 x 1 mL of 0.2% SDS in 1X PBS, 1X PBS, and $ddH_2O$). Enriched proteins were eluted via incubation in 3X Laemmli buffer with 25 mM biotin at 95 °C for 10 min. To remove beads, the tubes were centrifuged at 17,000 x g for 1 min, and the supernatant was transferred to another tube. The elution process was repeated with fresh buffer, and the combined supernatant was analyzed by silver staining and Western blot.

**Anaerobic Isolation of Gut Microbiota from Mouse Feces:**

Fecal pellets were weighed in sterile 15 mL conical tubes, and an equal weight of 3 mm glass beads were added. Tubes were flushed thoroughly with argon and moved into the Coy chamber on ice. Degassed, chilled 1X PBS (3 mL for 1 g feces) was then added, the tubes tightly capped and sealed with parafilm, and the feces were crushed by vortexing for 10 min at 4 °C. The slurry was then centrifuged at 300 x g for 5 min at 4 °C, with acceleration set to 3 and deceleration at 9. The supernatant was transferred to a 50 mL conical tube in the chamber. Degassed 1X PBS (same amount as above) was added to the pellets again, and the above sequence was repeated twice. Sterile, degassed Nycodenz solution (70% w/v, 500 µL) was added to 1.5 mL tubes in the chamber, and 1 mL of the supernatant was carefully layered on top. The biphasic solution was centrifuged at 10,000 x g for 40 min at 4 °C. After centrifugation, the top layer was carefully removed inside the chamber, leaving the middle layer of bacteria undisturbed. Then, the bacterial layer was removed, combined into a fresh 15 mL conical tube, mixed thoroughly, and aliquoted into clean 1.5 mL microfuge tubes. Bacteria were washed with 1X PBS (1 mL) three times, followed by centrifugation at 17,000 x g for 10 min at 4 °C. The final pellet was resuspended in 1X PBS containing 1 mM PMSF (300 µL for 150 µL bacterial pellets), and the samples were transferred to argon-flushed 2 mL screw cap tubes containing 330 µg of 0.1 mm zirconia/silica beads. The tubes were sealed with parafilm, then were subjected to agitation on a bead beater at 2,400 rpm (6 x 1 min, with 3 min rest on ice between). Finally, samples were centrifuged at 17,000 x g for 20 min at 4 °C, and the resulting supernatants were combined and transferred to a clean tube. Final protein concentration was quantified using the DC assay (Bio-Rad). Note: All steps aside from vortexing and centrifugation were carried out in an anaerobic chamber with degassed media. A minimal amount of ice was used to keep samples cool during isolation in the chamber, and no fluctuations in $O_2$ concentration was observed throughout the protocol.

**In-gel Fluorescence Assays with Mouse Fecal Bacteria:**

In the Coy chamber, lysates (100 µg) were prepared as described above and diluted to a final volume of 50 µL with 1X PBS for probe labeling (for CA-FMK reactions, NaOAc pH 5.5 was added for a final concentration of 50 mM and DTT was added for a final concentration of 1 mM). Lysates were treated with CA-FMK (100 µM), CA-Dimedone, or vehicle (DMSO) for 1 h at 37 °C in the chamber. After preclearing, the samples were precipitated, dried for 10 min at 37 °C, and resolubilized in 95 µL click buffer (0.1 M sodium phosphate buffer, 4% SDS, pH 7.4). Reagents were added into a master mix in the following final concentrations: 2 mM THPTA, 1 mM $CuSO_4$, 10 µM AZDye 647-alkyne, and 50 mM sodium ascorbate, in a final reaction volume of 120 µL. Click reactions were performed for 1 h at 37 °C then quenched via protein precipitation as previously described. Pellets were resuspended in 30 µL of 1X Laemmli buffer and boiled at 95 °C for 10 min, then analyzed by SDS-PAGE. Gels were destained (40% methanol, 50% acetic acid, 10% $ddH_2O$) on a rocker in the dark for 20 min, then visualized using a Bio-Rad ChemiDoc MP Imaging System. Lastly, gels were stained with Coomassie Brilliant Blue dye for 2 min, then destained (30% methanol, 5% acetic acid, 65% $ddH_2O$) overnight with gentle rocking prior to visualization using a Bio-Rad ChemiDoc MP Imaging System.

**Biotin-Streptavidin Enrichment of Labeled Protein from Mouse Fecal Bacteria:**

In the Coy chamber, lysates (1 mg) were prepared as described above and diluted to a final volume of 600 µL with 1X PBS for probe labeling (for CA-FMK reactions, NaOAc pH 5.5 was added for a final concentration of 50 mM and DTT was added for a final concentration of 1 mM). Lysates were treated with CA-FMK (100 µM), CA-Dimedone, or vehicle (DMSO) for 1 h at 37 °C in the chamber. Following probe treatment, lysates were diluted to 700 µL with 1.2% SDS in 1X PBS for preclearing. Streptavidin agarose beads (10 µL per 1 mg of lysate) were washed three times with 1 mL of 1X PBS containing 0.2% SDS, then added to the samples and incubated for 1 h with rotation. After preclearing, the samples were precipitated, dried for 10 min at 37 °C, and resolubilized in click buffer (0.2% SDS final concentration, 1X PBS, pH 7.4). Reagents were added into a master mix in the following final concentrations: 2 mM THPTA, 1 mM $CuSO_4$, 100 µM biotin-alkyne, and 50 mM sodium ascorbate, in a final reaction volume of 600 µL. Click reactions were performed for 1 h at 37 °C, then quenched via protein precipitation as previously described. After drying the pellets for 10 min at 37 °C, they were again resuspended in 1X PBS with a final concentration of 0.2% SDS. Streptavidin agarose beads (10 µL per 1 mg lysate) were washed three times with 1 mL 0.2% SDS in 1X PBS, then added to the samples and incubated for 1 h with rotation. After enrichment, tubes were centrifuged at 2,500 x g for 1 min at 4 °C, the supernatant was discarded, and the beads were then washed sequentially (3 x 1 mL of 0.2% SDS in 1X PBS, 1X PBS, and $ddH_2O$). Enriched proteins were eluted via incubation in 3X Laemmli buffer with 25 mM biotin at 95 °C for 10 min. To remove beads, the tubes were centrifuged at 17,000 x g for 1 min, and the supernatant was transferred to another tube. The elution process was repeated with fresh buffer, and the combined supernatant was analyzed by silver staining and Western blot.

**Silver Staining Analysis:**

The eluted proteins from biotin-streptavidin enrichment were analyzed by SDS-PAGE. First, the gels were fixed (50% MeOH, 5% acetic acid, 45% dd$H_2O$) for 20 min, then washed with 50% MeOH and dd$H_2O$ for 10 min each. The gels were then sensitized via incubation in 0.02% (w/v) sodium thiosulfate for 1 min, then washed with dd$H_2O$. Next, gels were incubated in 0.1% (w/v) silver nitrate with 0.08% paraformaldehyde for 20 min, then washed with dd$H_2O$ to remove excess silver. Gels were developed by adding 2% (w/v) sodium carbonate with 0.04% paraformaldehyde and gentle rocking until bands began to appear. After the desired level of signal to noise was obtained, 1-2 mL of acetic acid was added to quench the reaction. Finally, gels were washed with dd$H_2O$ and visualized using a Bio-Rad ChemiDoc MP Imaging System.

**Western Blot Analysis:**

The eluted proteins from biotin-streptavidin enrichment were analyzed by SDS-PAGE. After transfer to a nitrocellulose membrane using a turbo transfer system (Bio-Rad), the membrane was blocked with 5% BSA in TBST (1X tris-buffered saline with 0.1% Tween 20) for 1 h with gentle rocking, after which the membrane was incubated with streptavidin-HRP for 1 h with gentle rocking. Blots were then washed with TBST (3 x 5 min), developed with SuperSignal West Pico substrates (Thermo Fisher), and imaged using a Bio-Rad ChemiDoc MP Imaging System.

**In-gel Trypsin Digestion and Extraction of Enriched BSHs:**

Triplicate elutions of each probe or vehicle (DMSO, CA-Dim, CA-FMK) from streptavidin enrichment (30 μL) were loaded onto a 12% Bis/Tris SDS-PAGE gel and stained with SYPRO Ruby (Thermo Fisher). Each lane was excised separately between 37-50 kDa, cut into ~1 mm cubes, and subjected to in-gel digestion using trypsin. Briefly, the excised gel pieces were washed twice with dd$H_2O$ and 50 mM $NH_4HCO_3$ (50% acetonitrile (ACN):dd$H_2O$) and then were dehydrated with 100% ACN. Gel pieces were then reduced with 10 mM DTT (100 mM $NH_4HCO_3$ in dd$H_2O$; 150 μL) for 1 h at 55 °C, followed by alkylation with 55 mM iodoacetamide in the dark for 45 min at rt. Wash steps were then repeated as described above. Each sample was rehydrated with trypsin (Promega sequencing grade) at 1:10 (w/w) ratio in 50 mM $NH_4HCO_3$ (10% ACN:dd$H_2O$), followed by incubation at 37 °C for 16 h. The digested peptides were extracted twice with Buffer A (5% formic acid (FA), 50% ACN:dd$H_2O$), and once with Buffer B (5% FA, 90% ACN:dd$H_2O$). Supernatants from each extraction were combined, dried in a speed vacuum, reconstituted in 100 μL of dd$H_2O$, and filtered with a 0.22 μm spin filter (Costar Spin-X, Corning). Samples were then dried again, prior to being reconstituted into 2% ACN with 0.5% FA prior to LC-MS/MS analysis.

**Investigation of *C. perfringens* Cys2 Post-Translational Modifications using LC-MS/MS:**

Purified *C. perfringens* CGH (30 μg) was treated with $H_2O_2$ (1 mM) for 1 min, then with dimedone (100 μM) at 37 °C for 30 min. BSH was then precipitated as described above, resuspended in 30 μL 1X Laemmli Buffer, and subjected to SDS-PAGE.

In-gel trypsin digestion as described above was employed to obtain dried peptides of treated BSH, which were then reconstituted in 2% ACN containing 0.5% FA and 10 fmol/μL enolase (yeast) tryptic digest as an internal standard for nanoLC-ESI-MS/MS analysis. The analysis was carried out using an Orbitrap Fusion™ Tribrid™ (Thermo-Fisher Scientific, San Jose, CA) mass spectrometer equipped with a nanospray Flex Ion Source, and coupled with a Dionex UltiMate 3000 RSLCnano system (Thermo, Sunnyvale, CA)[4,5]. The peptide samples were injected onto a PepMap C-18 RP viper trapping column (5 µm, 100 µm i.d x 20 mm) at 20 µL/min flow rate for rapid sample loading and then separated on a PepMap C-18 RP nano column (2 µm, 75 µm x 25 cm) at 35 °C. The peptides were eluted in a 90 min gradient of 5% to 35% ACN in 0.1% FA at 300 nL/min, followed by an 8-min ramping to 90% ACN-0.1% FA and a 7-min hold at 90% ACN-0.1% FA. The column was re-equilibrated with 0.1% FA for 25 min prior to the next run. The Orbitrap Fusion was operated in positive ion mode with spray voltage set at 1.6 kV and source temperature at 275°C. External calibration for FT, IT and quadrupole mass analyzers was performed. In data-dependent acquisition (DDA) analysis, the instrument was operated using FT mass analyzer in MS scan to select precursor ions followed by 3 second "Top Speed" data-dependent CID ion trap MS/MS scans at 1.6 m/z quadrupole isolation for precursor peptides with multiple charged ions above a threshold ion count of 10,000 and normalized collision energy of 30%. MS survey scans at a resolving power of 120,000 (fwhm at m/z 200), for the mass range of m/z 375-1,600. Dynamic exclusion parameters were set at 35 s of exclusion duration with ±10 ppm exclusion mass width. All data were acquired under Xcalibur 4.4 operation software (Thermo-Fisher Scientific).

DDA RAW files were searched using Proteome Discoverer 3.1 software (Thermo Fisher Scientific, Bremen, Germany) using the *C. perfringens* database from UniProt (Proteome ID UP000000818). Precursor tolerance was set to 10 ppm and fragment mass tolerance was set to 0.5 Da. *In silico* digestion was performed with trypsin allowing for 4 missed cleavages. Two searches were performed: in the first, variable modifications searched included methionine oxidation, N-terminal acetylation, N-terminal methionine loss, cysteine carbamidomethylation, and sulfinic/sulfonic/dimedone modifications of cysteine. In the second search, the same variable modifications were used except cysteine sulfenic acid was substituted for sulfinic and sulfonic oxidation on cysteine. The searches are in separate tabs of the corresponding supplemental dataset file.

**Identification of Enriched Proteins from Anaerobes using LC-MS/MS Analysis:**

Dried tryptic digests from streptavidin enrichment from anaerobes as described above were reconstituted in 2% ACN containing 0.5% FA and 10 fmol/µL enolase (yeast) tryptic digest as an internal standard for nanoLC-ESI-MS/MS analysis. The analysis was carried out using an Orbitrap Fusion™ Tribrid™ (Thermo-Fisher Scientific, San Jose, CA) mass spectrometer equipped with a nanospray Flex Ion Source, and coupled with a Dionex UltiMate 3000 RSLCnano system (Thermo, Sunnyvale, CA)[4,5]. The peptide samples were injected onto a PepMap C-18 RP viper trapping column (5 µm, 100 µm i.d x 20 mm) at 20 µL/min flow rate for rapid sample loading and then separated on a PepMap C-18 RP nano column (2 µm, 75 µm x 25 cm) at 35 °C. The peptides were eluted in a 90 min gradient of 5% to 35% ACN in 0.1% FA at 300 nL/min, followed by an 8-min ramping to 90% ACN-0.1% FA and a 7-min hold at 90% ACN-0.1% FA. The column was re-equilibrated with 0.1% FA for 25 min prior to the next run. The Orbitrap Fusion was operated in positive ion mode with spray voltage set at 1.6 kV and source temperature at 275°C. External calibration for FT, IT and quadrupole mass analyzers was performed. In data-dependent acquisition (DDA) analysis, the instrument was operated using FT mass analyzer in MS scan to select precursor ions followed by 3 second "Top Speed" data-dependent CID ion trap MS/MS scans at 1.6 m/z quadrupole isolation for precursor peptides with multiple charged ions above a threshold ion count of 10,000 and normalized collision energy of 30%. MS survey scans at a resolving power of 120,000 (fwhm at m/z 200), for the mass range of m/z 375-1,600. Dynamic exclusion parameters were set at 35 s of exclusion duration with ±10 ppm exclusion mass width. All data were acquired under Xcalibur 4.4 operation software (Thermo-Fisher Scientific).

DDA RAW files were searched using MSFragger (v.4.1) and Fragpipe (v22) using *B. longum* ATCC 15697 and *B. fragilis* ATCC 25285 proteomes generated by predicted open reading frames using Prodigal[6–8]. Precursor and fragment mass tolerance was set to 20 ppm, *in silico* digestion was performed with strict trypsin allowing for 2 missed cleavages and peptide lengths of 5-30 amino acids, and peptide mass range set to 500-5000 kDa. Variable modifications searched included methionine oxidation, N-terminal acetylation, and N-terminal methionine loss. Cysteine carbamidomethylation was set as a static modification.

**Identification of Enriched BSHs from Mouse Fecal Bacteria using LC-MS/MS Analysis:**

The dried tryptic digests from each triplicate sample (9 samples total) were reconstituted in 20 µL of 0.5% FA for nanoLC-ESI-MS/MS analysis, which was carried out using an Orbitrap Eclipse™ Tribrid™ (Thermo-Fisher Scientific, San Jose, CA) mass spectrometer equipped with a nanospray Flex Ion Source, and coupled with a Dionex UltiMate3000RSLCnano system (Thermo, Sunnyvale, CA)[4,9]. The peptide samples (20 µL) were injected onto a PepMap C-18 RP nano trapping column (5 µm, 100 µm i.d x 20 mm) at 20 µL/min flow rate at 35 °C for rapid sample loading and then separated on an Aurora Ultimate C18 nano column (1.7 µm, 75 µm x 25 cm) at 50 °C using an IonOpticks system. The tryptic peptides were eluted in a 90 min gradient of 5% to 35% ACN in 0.1% FA at 300 nL/min, followed by a 7 min ramping to 90% ACN-0.1% FA and an 8 min hold at 90% ACN-0.1% FA. The column was re-equilibrated with 0.1% FA for 25 min prior to the next run. The Orbitrap Eclipse was operated in positive ion mode with spray voltage set at 1.25 kV and source temperature at 300°C. External calibration for FT, IT and quadrupole mass analyzers was performed. In data-dependent acquisition (DDA) analysis, the instrument was operated using FT mass analyzer in MS scan to select precursor ions followed by 3 second "Top Speed" data-dependent CID ion trap MS/MS scans at 1.6 m/z quadrupole isolation for precursor peptides with multiple charged ions above a threshold ion count of 10,000 and normalized collision energy of 32%. MS survey scans at a resolving power of 120,000 (fwhm at m/z 200), for the mass range of m/z 375-1,575. Dynamic exclusion parameters were set at 30 s of exclusion duration with ±10 ppm exclusion mass width. All data were acquired under Xcalibur 4.2 operation software (Thermo-Fisher Scientific).

RAW files were searched using MSFragger (v.4.1) and Fragpipe (v22)[6]. The default workflow settings for LFQ-MBR utilizing IonQuant (v 1.10.27)[10] were used with minimal modification as described below. Precursor and fragment mass tolerance was set to 20 ppm, *in silico* digestion was performed with strict trypsin allowing for 2 missed cleavages and peptide lengths of 5-30 amino acids, and peptide mass range set to 500-5,000 kDa. Due to the extremely large database size, mass calibration and optimization were turned off, and database splitting set to 4. Variable modifications searched included methionine oxidation, N-terminal acetylation, and N-terminal methionine loss. Cysteine carbamidomethylation was set as a static modification.

MSFragger results were filtered to remove contaminants and LFQ intensities were used to measure protein abundance and fold change, without normalization to a specific condition. Significantly enriched BSHs can be considered those with a FC ratio of greater than or equal to 2, and with p values smaller than 0.05.

**Metaproteomics Analysis:**

The metaproteomics database was constructed using the Mouse Gastrointestinal Bacteria Catalog (MGBC)[11]. Briefly, the nonredundant protein catalog was clustered to 95% identity using CD-HIT (v4.8.1)[12]. The clustered proteins were combined with the *Mus musculus* reference proteome (UniProt proteome ID: UP000000589, accessed July 17, 2025) and a set of common proteomics contaminants. Decoys were generated by protein sequence reversal[13]. The database was then searched using Fragpipe (v22.0).

**Annotation of Bile Salt Hydrolases:**

A previously curated set of microbiome BSHs[14] were aligned with MUSCLE (v5.1)[15] and used to generate a BSH-specific HMM using HMMer[16]. This HMM was then used to search the clustered MGBC protein catalog ($E$-value < $10^{-10}$). Sequences passing this threshold were then aligned with *C. perfringens* strain 13 BSH as a reference sequence using MUSCLE. From this alignment, candidate BSH sequences were identified based on the presence of conserved residues C2, R18, D21, N175, and R228 relative to *C. perfringens* strain 13 BSH[14].

**Phylogenetic Analysis of Bile Salt Hydrolases:**

Phylogenetic tree construction was performed in R (v 4.2.2). Briefly, BSH proteins identified by proteomics were aligned using Clustal Omega[17]. Distances were calculated using BLOSUM62[18] for neighbor-joining tree estimation[19,20] using phangorn[21]. Bootstrap values were calculated after performing $n$ = 100 resamples[22,23].

**Chemical Synthesis and Characterization**

**General Materials and Methods:**

Chemicals and reagents were purchased from a variety of vendors, including Sigma Aldrich, Ambeed, Acros, Fisher Scientific, Santa Cruz, AstaTech, Oakwood, TCI, Chem-Impex, Alfa Aesar, Click Chemistry Tools, Cayman, BroadPharm, VWR, and Cambridge Isotope Laboratories, and were used without further purification. Anhydrous solvents were obtained as commercially available pre-dried, oxygen-free formulations. Flash chromatography was carried out using Silicycle Siliaflash P60 40-60 Å silica gel. All reactions were monitored by thin layer chromatography (TLC) carried out on silica gel plates (60 Å) and visualized with UV light and ceric ammonium molybdate (CAM) stain. NMR spectra were recorded on Bruker AVIII-400 or Bruker AVIII- 500 in the indicated solvent. Multiplicities are reported with the following abbreviations: s singlet; d doublet; t triplet; m multiplet; br broad. Chemical shifts are reported in ppm relative to the residual solvent peak, and *J* values are reported in Hz. Reverse Phase HPLC separation was carried out on a Shimadzu system using a Luna Omega 5 µM 100 Å C18 analytical column (250 x 4.6 mm; flow rate 1 mL/min) and a Luna Omega 5 µM 100 Å C18 preparative column (250 x 21.2 mm; flow rate 20 mL/min). High Resolution Mass spectrometry data were collected on an Agilent 6500 Series Q-TOF LC-MS.

### Synthetic Procedures

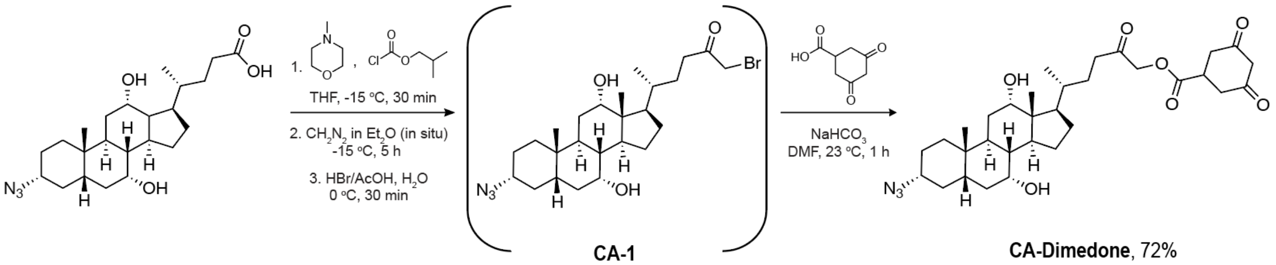

Scheme 1. Synthesis of CA-Dimedone

Azidocholic acid, CA1, and CA-FMK were synthesized as previously described[1,16].

Azidocholic acid (1.25 mmol) was dissolved in anhydrous THF (4.2 mL) and stirred at -15 °C for 10 min. Then, N-methylmorpholine (1.5 eq.) and isobutyl chloroformate (1.5 eq.) were sequentially added. The mixture was stirred at -15 °C for an additional 30 min, during which time a white precipitate formed. In the meantime, ethereal diazomethane was generated *in situ* according to the procedure reported in the Sigma Aldrich technical bulletin (AL-180). A flame polished glass pipette was used to add diazomethane (3.75 eq.) dropwise to the reaction mixture at 0 °C, then reaction was slowly warmed to room temperature over 5 h. To generate the corresponding bromomethyl ketone, hydrogen bromide (33 w% in acetic acid, 50 eq.) was mixed with 10 mL of dd$H_2O$ and added dropwise to the reaction mixture. The reaction was diluted and extracted with ethyl acetate, and then the organic layer was washed with dd$H_2O$, $NaHCO_3$, and brine. After drying over anhydrous $Na_2SO_4$, the organic layer was further concentrated *in vacuo* to yield a sticky yellow solid. The crude **CA-1** was used directly without further purification.

Safety Note: Significant hazards were mitigated in the generation of diazomethane by using ground glass joints, a blast shield, and loosely closing the reaction vessel. All syringes, needles, and glassware used in the reaction were quenched with acetic acid prior to disposal or cleaning.

**Characterization of Probes:**

**CA-Dimedone (CA-Dim):**

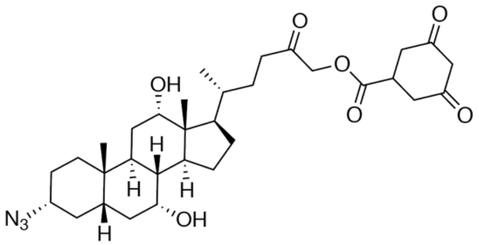

Crude **CA-1** was dissolved in anhydrous DMF (2.5 mL), then 3,5-dioxocyclohexane-1-carboxylic acid (1 eq.) and sodium bicarbonate (1.1 eq.) were added, and the reaction was stirred under an inert atmosphere for 2 h at 25 °C. The mixture was diluted and extracted with ethyl acetate, dried over anhydrous $Na_2SO_4$, and further concentrated *in vacuo*. The crude was purified by flash chromatography (20→100% of ethyl acetate in hexanes) and preparative HPLC (20→100% of ACN in $H_2O$ over 30 min) to yield a sticky white solid (72% yield). $^1$H NMR (500 MHz, MeOD) δ 3.95 (br s, 1H), 3.81(br s, 1H), 3.65 (s, 1H), 3.2-3.12 (m, 1H), 2.67-2.6 (m, 1H), 2.56-2.49 (m, 1H), 2.44-2.35 (m, 2H), 2.28-2.19 (m, 2H), 2.04-1.94 (m, 3H), 1.92-1.83 (m, 4H), 1.8-1.73 (m, 2H), 1.69-1.63 (m, 2H), 1.6-1.48 (m, 7H), 1.46-1.39 (m, 3H), 1.37-1.26 (m, 3H), 1.16-1.09 (m, 1H), 1.05-1.03 (m, 1H), 1.00 (d, J = 6.5 Hz, 3H), 0.94 (s, 3H), 0.71 (s, 3H). $^{13}$C NMR (126 MHz, MeOD) δ 205.13, 205.09, 205.04, 176.53, 73.96, 68.86, 62.74, 52.00, 47.94, 47.47, 43.41, 42.98, 40.97, 37.28, 36.71, 36.64, 36.60, 35.95, 35.66, 32.22, 31.83, 30.71, 30.69, 29.54, 28.65, 27.86, 27.79, 24.19, 23.14, 17.73, 17.56, 12.97. HRMS (ESI-TOF-MS): $[M+NH_4]^+$ calcd. for $C32H51N4O7^+$ 603.3753; found 603.3778.

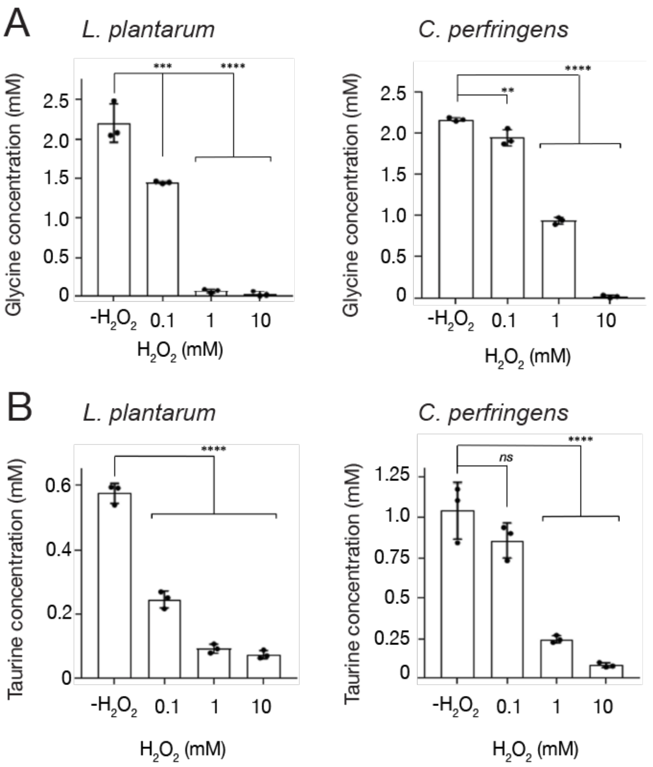

**Figure S1. BSH activity assays show oxidation inhibits product formation.** (A) Activity assays measuring the extent of deconjugation of glycocholate by purified CGH (from *C. perfringens*) and BSH (from *L. plantarum*) with and without $H_2O_2$ exposure for 1 min at the indicated concentrations. Concentration of free glycine was used as the measure of enzyme activity. (B) Activity assays measuring the extent of deconjugation of taurocholate by purified CGH (from *C. perfringens*) and BSH (from *L. plantarum*) with and without $H_2O_2$ exposure for 1 min at the indicated concentrations. Concentration of free glycine was used as the measure of enzyme activity. Error bars represent standard deviation from the mean. One-way ANOVA, followed by post hoc Tukey's test: *** $p<0.001$, **** $p<0.0001$, n.s. = not significant, n = 3.

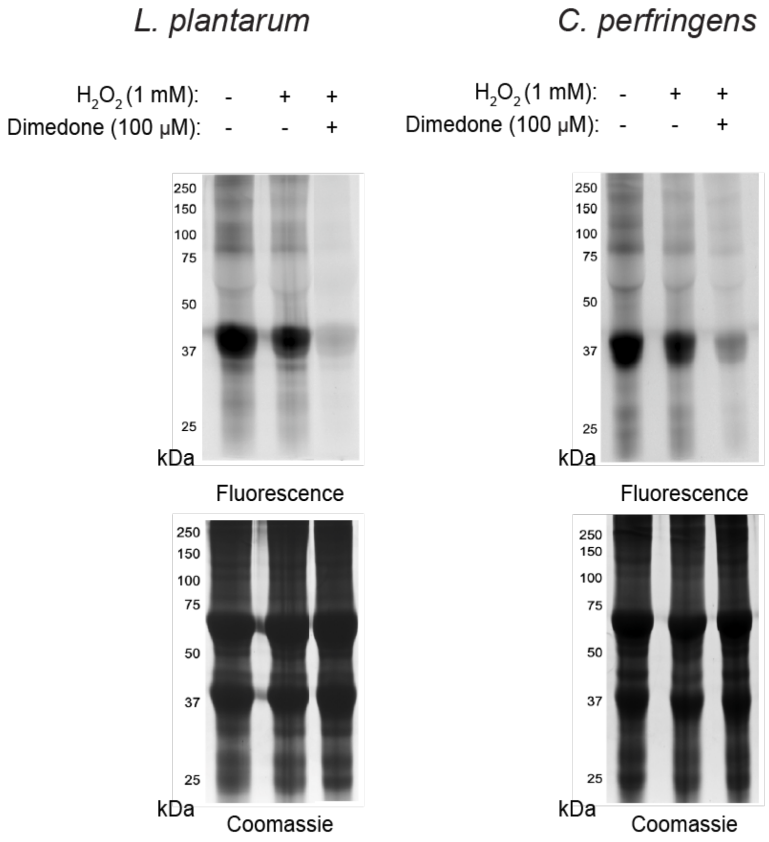

**Figure S2. Dimedone traps BSH Cys2 sulfenic acid formed after $H_2O_2$ addition.** In-gel fluorescence results from purified BSH from *L. plantarum* (left) and CGH from *C. perfringens* (right) that was treated with 1 mM $H_2O_2$ or vehicle (dd$H_2O$), then with dimedone (100 µM) for 30 min at 37 °C, lastly followed by CA-Dim or CA-FMK (10 µM) for 30 min at 37 °C. Coomassie shown to demonstrate equal protein loading.

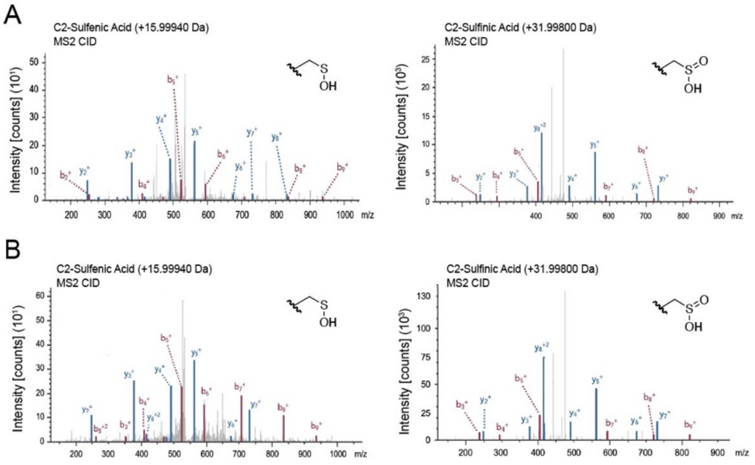

**Figure S3. LC-MS/MS analysis of *C. perfringens* CGH reveals oxidative PTMs on Cys2.** *C. perfringens* CGH was incubated with $H_2O_2$ (1 mM), followed by with or without dimedone (100

μM) for 30 min prior to LC-MS/MS analysis. (A) Representative MS/MS ($MS^2$) spectra showing detected oxPTMs on the Cys2 peptide from CGH treated with $H_2O_2$ only. Both the sulfenic (Cys-SOH, *left*) and sulfinic (Cys-$SO_2$H, *right*) acid states of Cys2 were present. (B) Representative $MS^2$ spectra showing detected oxPTMs on the Cys2 peptide from CGH treated with $H_2O_2$ and dimedone. Both the sulfenic (Cys-SOH, *left*) and sulfinic (Cys-$SO_2$H, *right*) acid states of Cys2 were present.

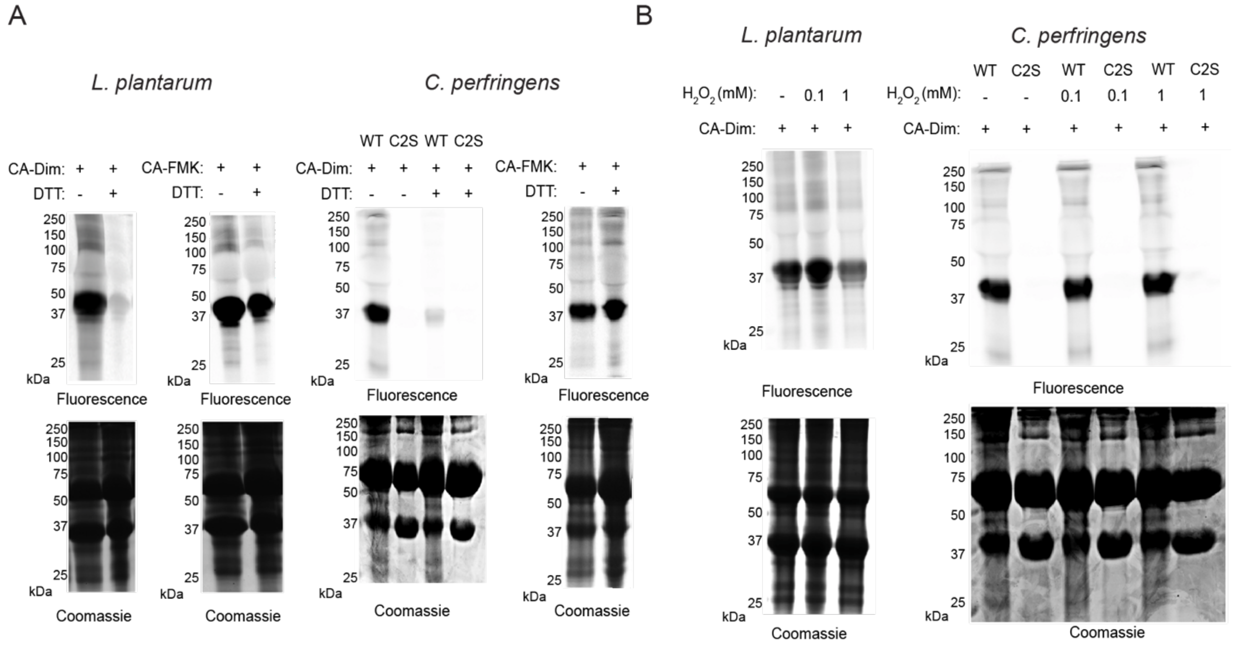

**Figure S4. Redox reagents modulate labeling of BSH by chemical probes.** (A) In-gel fluorescence results from purified BSH from *L. plantarum* (left) and CGH from *C. perfringens* (right) that were treated with DTT (20 mM) for 20 min at 55 °C, followed by CA-Dim or CA-FMK (10 μM) for 30 min at 37 °C. (B) In-gel fluorescence results from purified BSH from *L. plantarum* (left) and CGH from *C. perfringens* (right) that were treated with indicated concentrations of $H_2O_2$ for 1 min, followed by CA-Dim or CA-FMK (10 μM) for 30 min at 37 °C. (A-B) For *C. perfringens* CGH, the C2S mutant was included to show site-selectivity of CA-Dim probe labeling. After CuAAC tagging with AZDye 647-alkyne, labeling of BSH was visualized via in-gel fluorescence after SDS-PAGE. Coomassie shown to demonstrate equal protein loading.

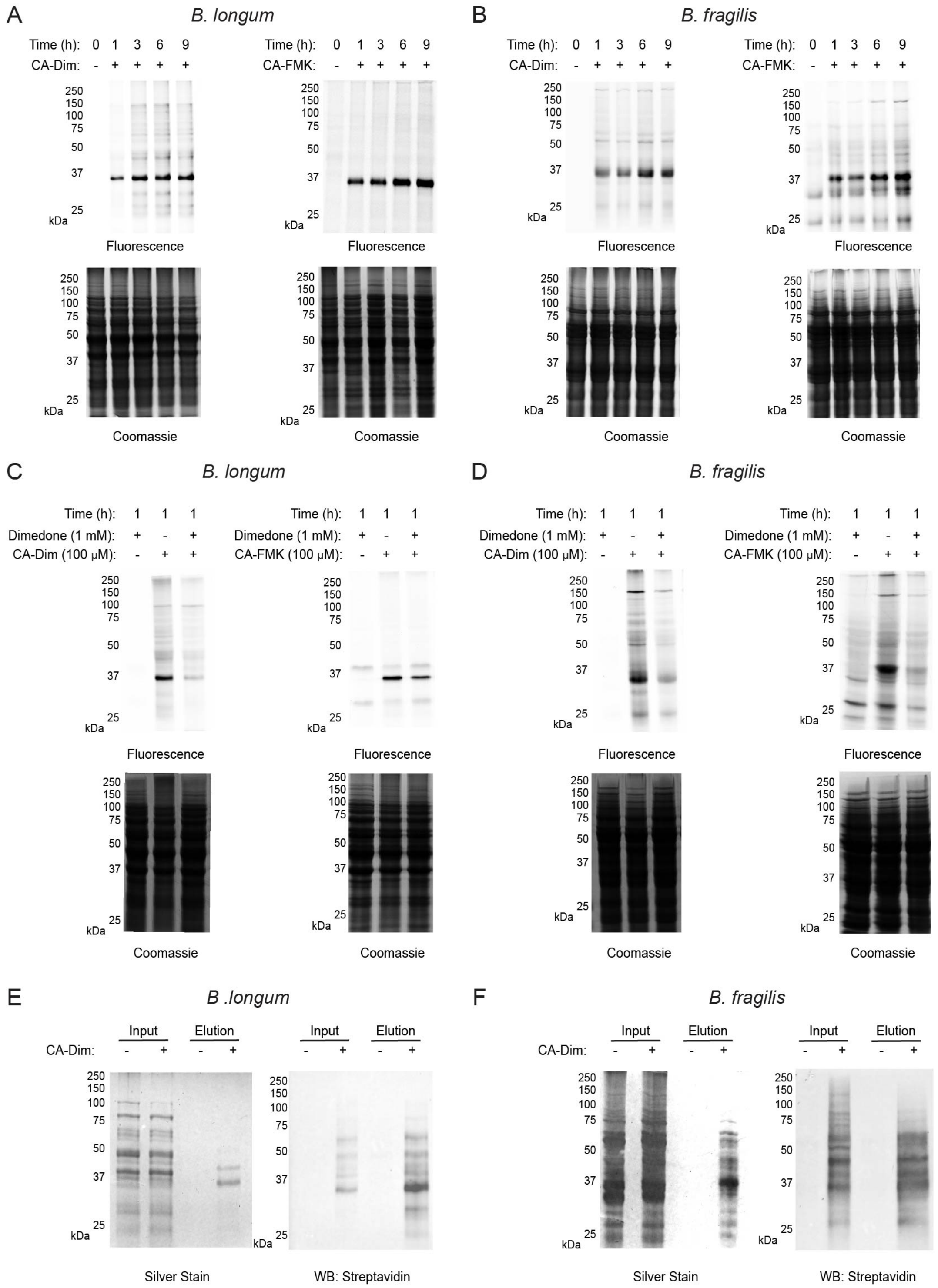
A
B. longum
B
B. fragilis
Time (h): 0 1 3 6 9
CA-Dim: - + + + +
CA-FMK: - + + + +
250
150
100
75
50
37
25
kDa
Fluorescence
Coomassie
C
B. longum
D
B. fragilis
Time (h): 1 1 1
Dimedone (1 mM): + - +
CA-Dim (100 µM): - + +
CA-FMK (100 µM): - + +
E
B .longum
F
B. fragilis
Input
Elution
Silver Stain
WB: Streptavidin

**Figure S5. Chemical probes label live anaerobes and demonstrate endogenous sulfenic acid formation**. (A) *B. longum* and (B) *B. fragilis* were treated with CA-Dim (*left*) or CA-FMK (*right*) at 100 µM for the indicated amount of time. (C) *B. longum* and (D) *B. fragilis* were treated with dimedone (1 mM) for 1 h, followed by CA-Dim (*left*) or CA-FMK (*right*) at 100 µM for 1 h. (E) *B. longum* and (F) *B. fragilis* were treated with CA-Dim (100 µM) for 12 h. After harvesting and lysing bacteria, CuAAC tagging with AZDye 647-alkyne or biotin-alkyne were performed, and labeling of BSH was visualized via in-gel fluorescence, silver stain, or Western blot via streptavidin-HRP after SDS-PAGE. Coomassie shown to demonstrate equal protein loading.

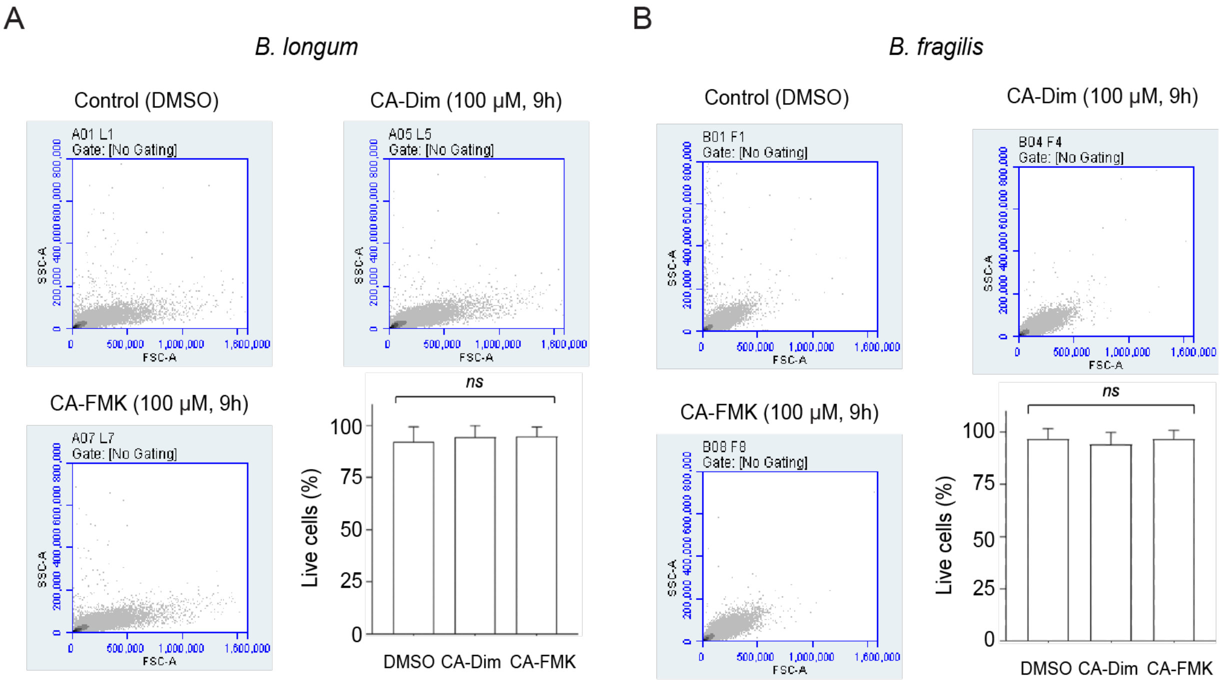

**Figure S6. Propidium iodide staining indicates probes are non-toxic to anaerobes.** Representative forward- versus side-scatter plots of (A) *B. longum* and (B) *B. fragilis* after live-labeling with CA-Dim or CA-FMK (100 µM) for 9 h. Percentage of live cells were plotted with normalization to the control. One-way ANOVA, followed by post hoc Tukey's test: n.s. = not significant, n = 3.

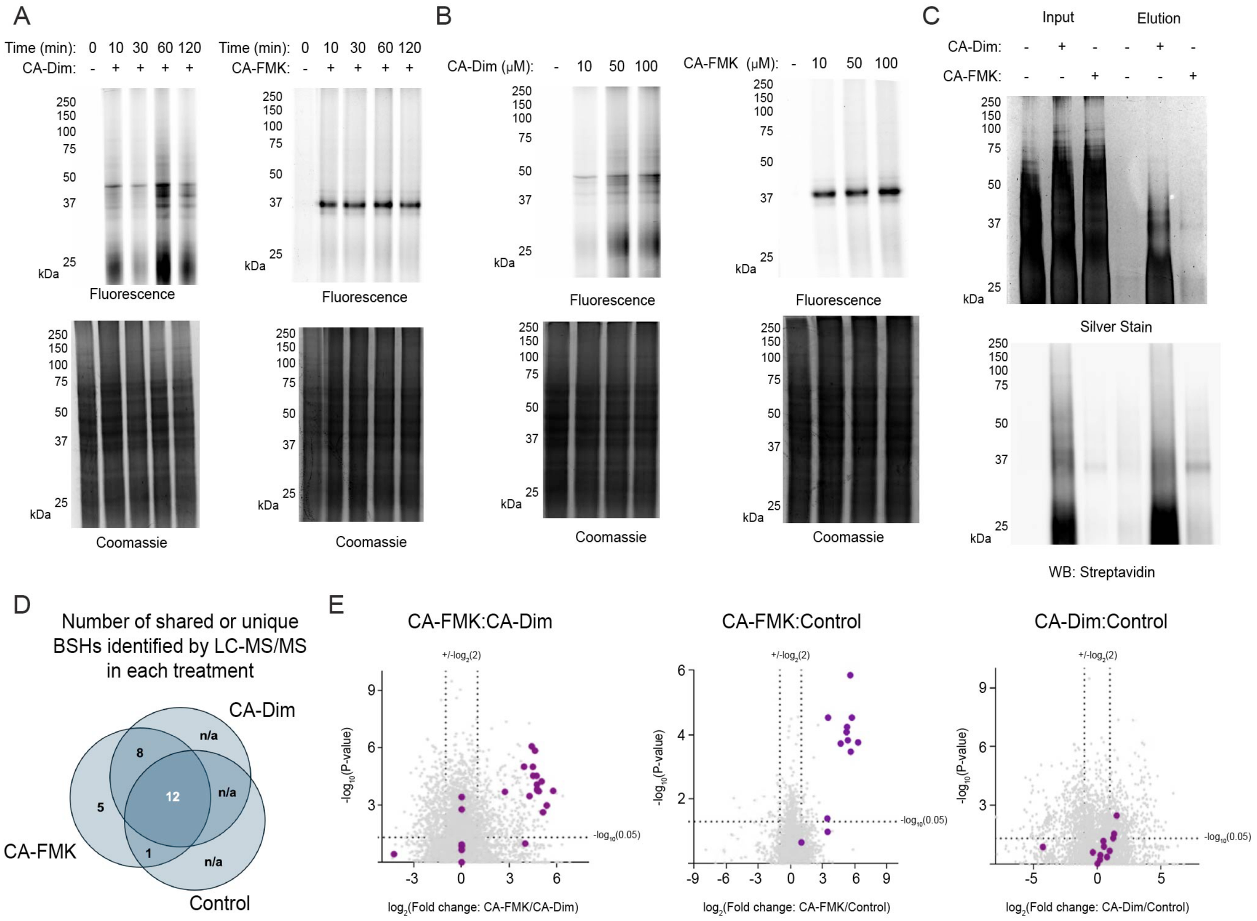

**Figure S7. Chemical probes label BSH in the murine gut microbiome.** Bacteria isolated anaerobically from fecal samples of wildtype mice were lysed and treated with CA-Dim (*left*) or CA-FMK (*right*) at 100 μM for the indicated amount of time (A), with varying concentrations at 1 h (B), or with both probes at 100 μM for 1 h (C). Then, CuAAC tagging with AZDye 647-alkyne or biotin-alkyne were performed, and labeling of BSH was visualized via in-gel fluorescence, silver stain, or Western blot via streptavidin-HRP after SDS-PAGE. Coomassie shown to demonstrate equal protein loading. (D) Venn diagram depicting number of BSHs identified by LC-MS/MS. (E) Volcano plots of indicated ratios from the proteomics results. Purple dots are BSH proteins.

$^{1}H$-NMR spectrum (500 MHz) of **CA-Dim** in $CD_3OD$

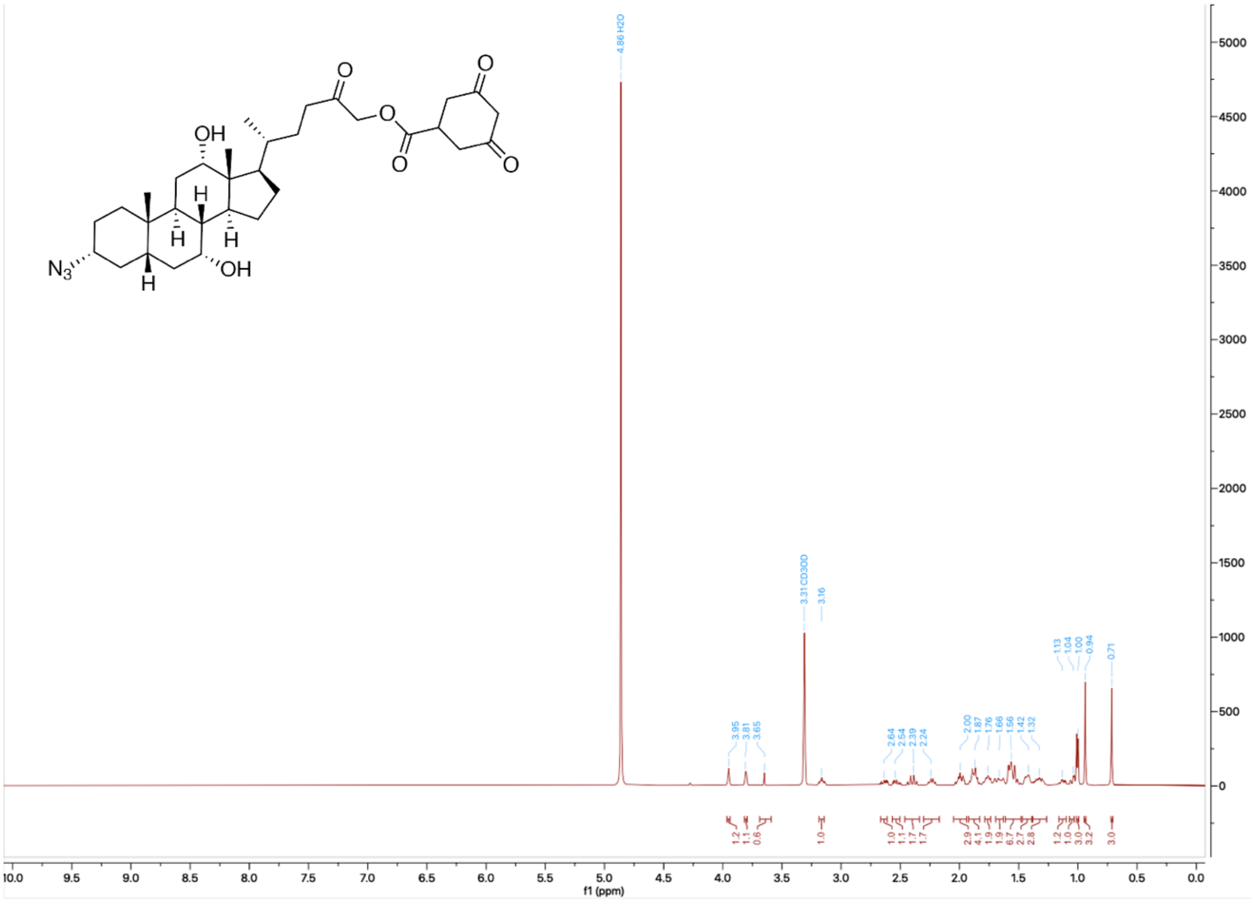

$^{13}$C-NMR spectrum (126 MHz) of **CA-Dim** in $CD_3OD$

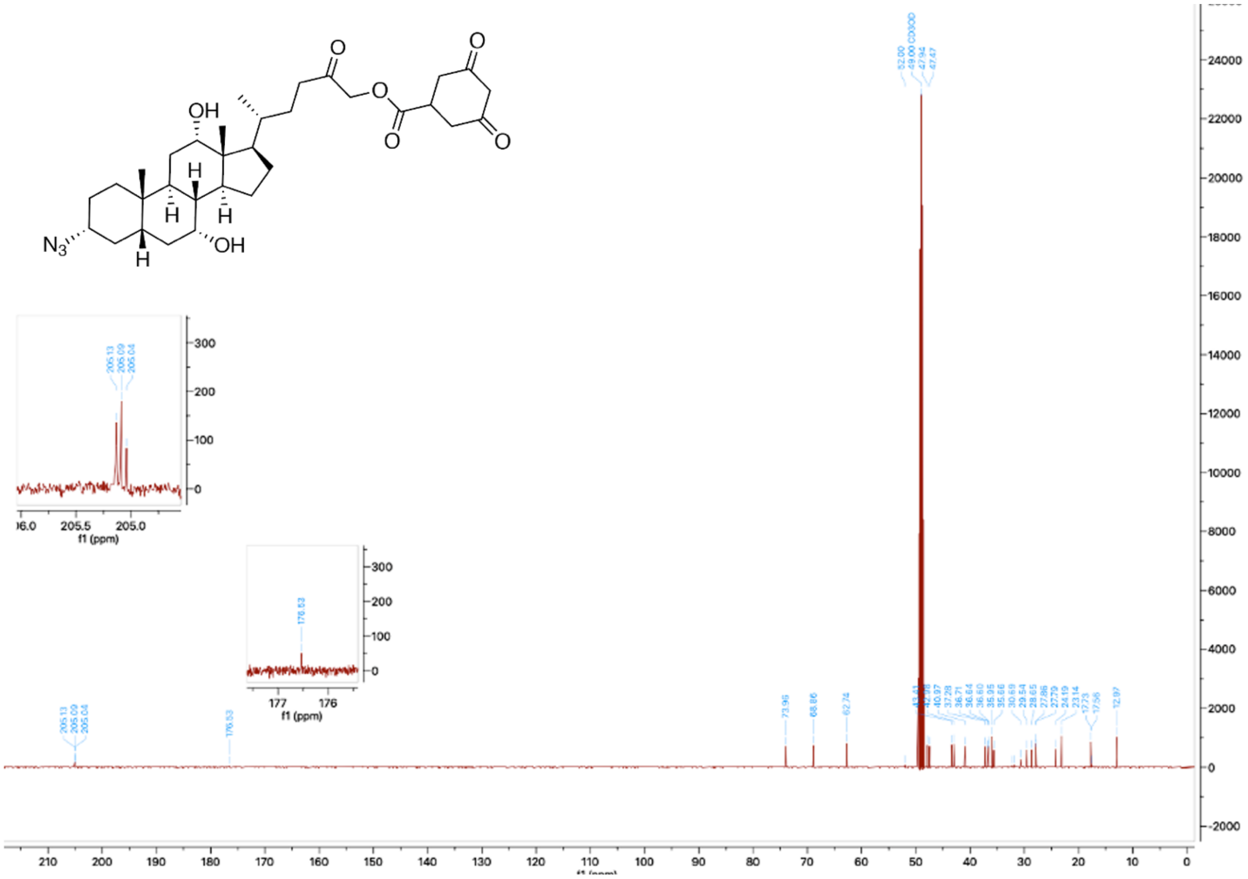
